# An integrated strategy reveals complex glycosylation of erythropoietin using top-down and bottom-up mass spectrometry

**DOI:** 10.1101/2021.02.09.430553

**Authors:** Yudong Guan, Min Zhang, Manasi Gaikwad, Hannah Voss, Ramin Fazel, Samira Ansari, Huali Shen, Jigang Wang, Hartmut Schlüter

## Abstract

The characterization of glycoproteins, like erythropoietin, is challenging due to the structural micro- and macro-heterogeneity of the protein glycosylation. This study presents an in-depth strategy for glycosylation analysis of a first-generation erythropoietin (epoetin beta), including a developed top-down mass spectrometric workflow for N-glycan analysis, bottom-up mass spectrometric methods for site-specific N-glycosylation and a LC-MS approach for O-glycan identification. Permethylated N-glycans, peptides and enriched glycopeptides of erythropoietin were analyzed by nanoLC-MS/MS and de-N-glycosylated erythropoietin was measured by LC-MS, enabling the qualitative and quantitative analysis of glycosylation and different glycan modifications (e.g., phosphorylation and O-acetylation). Extending the coverage of our newly developed Python script to phosphorylated N-glycans enabled the identification of 140 N-glycan compositions (237 N-glycan structures) from erythropoietin. The site-specificity of N-glycans was revealed at glycopeptide level by pGlyco software using different proteases. In total, 215 N-glycan compositions were identified from N-glycan and glycopeptide analysis. Moreover, LC-MS analysis of de-N-glycosylated erythropoietin species identified two different O-glycan compositions, based on the mass shifts between non-O-glycosylated and O-glycosylated species. This integrated strategy allows the in-depth glycosylation analysis of a therapeutic glycoprotein to understand its pharmacological properties and improving the manufacturing processes.

## INTRODUCTION

Erythropoietin (EPO) is renal growth factor that stimulates the proliferation and differentiation of erythroid-progenitor cells in the bone marrow and that is therefore used for the treatment of a wide spectrum of erythrocyte deficiency, which was commercialized in 1989 by Amgen as Epogen^®^.^1^ The binding of EPO to its receptor (EPOR), initiating the intracellular signaling pathways, highly depends on its amino acid sequence and post-translational modifications (PTMs), such as glycosylation and glycan modifications.^2^ EPO is available in four different forms, associated with different glycosylation patterns, including first-generation EPOs (epoetin alfa, epoetin beta and epoetin delta) and second-generation EPO (darbepoetin alfa). Darbepoetin alfa has five modified amino acids and two additional N-glycosylation sites compared to first-generation EPOs.^3^ First-generation epoetin consists of 166 amino acids with mono-, bi-, tri-, or tetra-antennary glycans structures on three N-glycosylation sites (Asn-24, −38, and −83) and one O-glycosylation site (Ser-126).

Glycosylation has significant impacts on the biological functions of a protein, including protein stability, solubility, antigenicity, folding and half-life.^4^ An in-depth characterization of glycan patterns is important to guarantee the proper manufacturing of biotherapeutic glycoproteins with appropriate pharmacological characteristics. In contrast to polypeptides, the biosynthesis of glycans is not directly depending on the templates, resulting in a high heterogeneity of glycoproteins in biotechnological productions. It has been shown that the glycan composition of the biopharmaceuticals is critically depending on the presence and activity of different glycosyltransferases, concentration of substrates, cell lines and cell culture conditions, making the production of therapeutic glycoproteins challenging.^5^ EPO is highly heterogeneous with a variety of potential glycosylation patterns as well as post-glycosylational modifications, such as sialylation by N-acetylneuraminic acid (Neu5Ac) or N-glycolylneuraminic acid (Neu5Gc), phosphorylation and O-acetylation.^6^ These patterns have a significant impact on the drug efficiency and adverse drug effects in EPO-treated patients. As an example, increases of the Neu5Gc containing glycoproteins can lead to immunogenic or inflammatory responses in patients.^7,8^ Phosphate groups are commonly attached on C6 position of mannose (Man) of N-glycans.^9,10^ O-acetyl groups bind to either C4-, C7-, C8-, or C9-hydroxyl position of sialic acid residues.^11^ For N-glycan sample preparation, O-acetylation is labile and removed during the commonly used permethylation, simplifying data analysis for N-glycan compositions.^12^ However, previous researches mainly analyzed EPO with total attached glycan compositions.^13-15^

Liquid chromatography (LC) coupled with electrospray ionization (ESI)-mass spectrometry (MS) and matrix-assisted laser desorption ionization (MALDI)-MS are used frequently to analyze biomolecules with an outstanding accuracy and reproducibility, like proteomics.^16-18^ For glycan analysis, various LC-MS-based strategies have been utilized, such as porous graphitic carbon (PGC)-LC-MS,^19^ hydrophilic interaction liquid chromatography (HILIC)-LC-MS^20,21^ and reversed-phase (RP)-LC-MS.^22^ In contrast to proteomics, MS-based glycomics is limited by differing standard procedures across different laboratories, technical limitations as well as the low availability of reliable glycoinformatics approaches, requiring efficient automated data interpretation software.^23^ NanoLC-MS/MS is more able to identify low abundant glycans than MALDI-MS, due to the ionization suppression during the laser desorption of MALDI-MS without chromatographic separation. An improved strategy to perform glycan-based permethylation and in-depth N-glycomics, using nanoLC(RP)-MS/MS combined with the developed R-script matching algorithm, significantly increases the coverage of identifiable N-glycans and has been reported in our previous study.^24^

Aiming to determine each glycan precisely unlike the previous researches that mainly analyzed EPO with total attached glycan compositions, an integrated analytical workflow is developed to analyze different glycosylation patterns of EPO (epoetin beta), expressed by Chinese hamster ovary (CHO) cell lines containing a cloned human erythropoietin gene, at N-glycan, glycopeptide and de-N-glycosylated protein levels. In this platform, N-glycans were measured by top-down approach, using nanoLC(RP)-MS/MS after N-glycan release, purification, reduction and permethylation, followed by data analysis with newly designed Python scripts. In addition, enriched glycopeptides were analyzed by bottom-up approach, using nanoLC(RP)-ESI-MS/MS after tryptic or chymotryptic digestion. LC-MS analysis of de-N-glycosylated EPO was performed, enabling the identification of O-glycans based on mass shift between non-O-glycosylated and O-glycosylated species. The glycosylation analysis of EPO at multiple levels provides new insights to profile the glycan structures, understand its pharmacological properties and control the manufacturing processes.

## MATERIALS AND METHODS

### Materials

All used chemicals, including sodium hydroxide beads, dimethylsulfoxide (DMSO) and iodomethane, were purchased from Sigma-Aldrich (Darmstadt, Germany), unless otherwise stated. A2F N-glycan standards were obtained from QA-Bio. Two different batches of EPOs (epoetin beta: RDF9729003 and RDF9729004) were kindly provided by CinnaGen (Tehran, Iran), which were herein named EPO-3 and EPO-4. Sequencing grade modified trypsin, chymotrypsin and peptide-N-glycosidase F (PNGase F) were obtained from Promega (Madison, WI, USA). Centrifugal filters (0.5 mL, 3 k) were purchased from Merck KGaA (Darmstadt, Germany). Solid-phase-extraction (SPE) columns containing reversed-phase (RP) materials (C_18_ Sep-Pak cartridges) were obtained from Waters (Miford, MA, USA).

### N-glycan release, purification, reduction and permethylation

N-glycan cleavage, purification, reduction and permethylation were performed as described in previous studies.^24,25^ EPO was denatured using 6 M urea, reduced by 30 mM dithiothreitol (DTT) at 56□ for 30 min and alkylated by 40 mM iodoacetamide (IAA) at room temperature for 30 min in dark, in which DTT and IAA were prepared in 100 mM ammonium bicarbonate (ABC) buffer. Next, buffers were exchanged to 100 mM ABC buffer using 3 kDa cut-off centrifugal filter. Thirty unit of PNGase F was added and incubated at 37°C for 24 h, followed by the addition of trypsin (1/100, w/w) for another 20 h. N-glycans and tryptic peptides were separated using a RP-SPE C_18_ cartridge. Briefly, the C_18_ cartridge was conditioned with 5 mL methanol and equilibrated with 10 mL 5% acetic acid respectively. After loading digested samples, N-glycans were eluted with 5 mL 5% acetic acid and evaporated using a SpeedVac™ vacuum concentrator. The A2F N-glycan standards, containing Neu5Ac_2_HexNAc_3_Hex_5_Fuc_1_Red-HexNAc_1_ at a purity of more than 90%, is collected from porcine thyroglobulin by hydrazinolysis using a combination of HPLC and glycosidase digestion. EPO derived N-glycans and A2F N-glycan standards were reduced by borane-ammonia complex, followed by optimized solid-phase permethylation (OSPP) with iodomethane. Briefly, 10 µg/µL borane-ammonia complex was used to reduce N-glycans at 60□ for 1 h and then removed by evaporation with three additions of 300 µL methanol. Reduced N-glycans were resuspended in 110 µL water/DMSO (10/100, v/v) solution and 100 µL iodomethane was added, which was transferred to the glass vial containing sodium hydroxide beads (200 mg). Afterwards, samples were incubated in a Thermomixer compact (Eppendorf AG, Hamburg, Germany) using a rotation speed of 1,300 rpm for 10 min. The solution containing permethylated N-glycans was transferred to a new vial, followed by the addition of 200 µL 5% acetic acid and 400 µL chloroform. Chloroform-water extraction was performed to purify permethylated N-glycans, which was dried by a SpeedVac™ vacuum concentrator and stored at −20°C until further use.

### NanoLC-MS/MS analysis of permethylated N-glycans

The permethylated N-glycans, resuspended into 0.1% formic acid (FA), were injected into a Dionex Ultimate 3000 UPLC system (Thermo Fisher Scientific, Bremen, Germany), coupled with a tribrid quadrupole-orbitrap-ion trap mass spectrometer (Fusion, Thermo Fisher Scientific, Bremen, Germany).^26^ Solvent A was 0.1% FA and solvent B was 0.1% FA in ACN. The sample was desalted using a RP C_18_ trapping column (Thermo Scientific™ Acclaim PepMap™, 100 μm×2 cm, 5 μm, 100Å) at a flow rate of 3 µL/min with 2% solvent B, and then transported to an analytical RP C_18_ column (Thermo Scientific™ Acclaim PepMap™ RSLC, 75 μm×50 cm, 2 μm, 100Å) at a flow rate of 0.2 µL/min. For N-glycan separation, solvent B started with 10%, increasing to 30% in 5 min, to 75% in 70 min and finally to 95% in 80 min. Detailed MS instrument parameters are described in Supporting Information. The data were visualized using Thermo Xcalibur in the version 4.2.28.14 (Thermo Fisher Scientific, Bremen, Germany).

### Identification and statistical analysis of permethylated N-glycans

For N-glycan identification, permethylated Neu5Gc, Neu5Ac, N-acetylhexosamine (HexNAc), Hexose (Hex), Fucose (Fuc) and reduced N-acetylhexosamine (Red-HexNAc) are utilized as the building blocks to match experimental masses with theoretical N-glycan compositions using a newly designed Python script glycoinformatics algorithm. With upgrading the previous R-scripts we have developed (https://github.com/guan181992/Glycoinformatics),^24^ which matched one precursor to at most one possible N-glycan composition within a predefined deviation, the newly designed Python script can provide all the possible N-glycan compositions for one precursor without N-glycan libraries, especially solving the problems derived from isobaric compositions like Neu5Ac_1_Hex_1_ and Neu5Gc_1_Fuc_1_ (Figure S1). It further extends to the N-glycans assembled with phosphorylated Man residues (P-Hex), considering the HexNAc_1_Hex_3_Red-HexNAc_1_ (trimannosylchitobiose core) (Figure S2). Mass spectrometric raw data was processed using the MaxQuant software in the version 1.6.2.3 (http://www.maxquant.org). As MaxQuant was exclusively used for mass value extraction, no database was included and all settings were chosen as automatically recommended for mass spectra.^27^ All the masses, in the resulting “allPeptides.txt” file, were input into Python script and matched with possible N-glycan compositions with a deviation threshold of 2.5 p.p.m. at MS1 level. From the result table, the masses, matched to N-glycan compositions, were further matched with monoisotopic m/z in the MS raw data. Finally, the positively matched masses were characterized as N-glycan structure with the assistance of GlycoWorkbench in the version 2.1 build 146 (https://download.cnet.com/GlycoWorkbench64-bit/3000-2383_4-75758804.html).

To perform quantitative analysis of N-glycans from different batches of EPO samples, EPO-3 and EPO-4, peak area of each identified N-glycan composition was integrated using Skyline in the version 19.1 (http://skyline.maccosslab.org). N-glycan compositions and Total Area MS1 (abundance of each N-glycan composition) were uploaded to Perseus in the version 1.6.2.1 (http://www.perseus-framework.org) for statistical analysis. Log2 transformation, normalization and two-sample student’s T-test at a false discovery rate (FDR) less than 0.05 were performed to identify significantly differential abundant N-glycans between the compared groups. Only N-glycans, showing a mean difference higher than 1.5 or 2-fold change (FC) between different EPO batches, were considered in further analysis.

### Tryptic and chymotryptic digestion of EPO

EPO samples were denatured, reduced and alkylated as described above. Buffers were exchanged to 100 mM ABC buffer using 3 kDa cut-off centrifugal filters. Samples were digested using trypsin (1/100, w/w) or chymotrypsin (1/100, w/w) at 37□ for 20 h, respectively. Resulting peptides were divided into two aliquots for peptide analysis and glycopeptide enrichment, which were concentrated by a SpeedVac™ vacuum concentrator and stored at −20□ until further use.

### NanoLC-MS/MS analysis and raw data processing for peptides

Digested EPO, by trypsin or chymotrypsin, was resuspended in 0.1% FA for nanoLC-MS/MS analysis. Solvent A was 0.1% FA and solvent B was 0.1% FA in ACN. The peptides were desalted using a RP C_18_ trapping column at a flow rate of 3 µL/min with 2% solvent B and then transported to an analytical RP C_18_ column at a flow rate of 0.2 µL/min. Starting with 1% for 5 min, solvent B increased to 25% in 65 min, to 35% in 80 min and finally to 90% in 85 min. Peptide ions were transferred to a hybrid quadrupole-orbitrap mass spectrometer (Q Exactive, Thermo Fisher Scientific, Bremen, Germany). Detailed MS instrument parameters are described in Supporting Information.

The peptide identification was performed using the pFind software in the version 3.1.6 (http://pfind.ict.ac.cn/software/pFind3/),^28^ with the FASTA database containing amino acid sequence of epoetin beta (https://www.drugbank.ca/drugs/DB00016). For peptide identification, up to 3 missed cleavages were tolerated; peptides identified at FDR less than 0.01 with a minimum length of 6 amino acids and a maximum length of 100 were considered; the precursor mass tolerance was set to 10 p.p.m.; the mass tolerance at MS2 was set to 20 p.p.m.; the oxidation of methionine residues was included as a variable modification; carbamidomethylation of cysteine residues was set as a fixed modification.

### Glycopeptide enrichment by zwitterionic hydrophilic interaction liquid chromatography

Digested EPO, by trypsin or chymotrypsin, was resuspended in 80% ACN with 1% trifluoroacetic acid (TFA) and enriched using self-packed zwitterionic hydrophilic interaction liquid chromatography (ZIC-HILIC) columns.^29^ Micro-columns were packed with 30 mg ZIC-HILIC materials obtained from a HILIC column (SeQuant ZIC-cHILIC, 4.6×250 mm, 3µm, 100Å, Merck KGaA, Darmstadt, Germany) on top of a C_8_ RP chromatographic materials (3M, Eagan, MN, USA). The column was equilibrated with 600 μL of 1% TFA in 80% ACN. After sample loading, low-hydrophilic peptides were removed using 600 µL of 1% TFA in 80% ACN. Glycopeptides were eluted by 600 µL 0.1 % TFA, concentrated by a SpeedVac™ vacuum concentrator and stored at −20□ until further use.

### NanoLC-MS/MS measurement and pGlyco-based data analysis of enriched glycopeptides

Dried glycopeptides were dissolved in 0.1% FA and injected into a Dionex Ultimate 3000 UPLC system, coupled to an orbitrap Fusion mass spectrometer. Solvent A was 0.1% FA and solvent B was 0.1% FA in ACN. Samples were firstly desalted using a RP C_18_ trapping column at a flow rate of 3 µL/min with 1% solvent B and then transported to an analytical RP C_18_ column at a flow rate of 0.2 µL/min. Solvent B started with 1% and increased to 20% in 90 min, to 30% in 120 min and to 90% in 130 min. Detailed MS instrument parameters are described in Supporting Information.

Glycopeptides were identified by pGlyco software in the version 2.2.0 (http://pfind.ict.ac.cn/software/pGlyco/index.html) with the total N-glycan entries of 7,884.^30^ By pGlyco identification, the amino acid sequence of epoetin beta was used as a FASTA database. The parameters were set as following: the precursor tolerance was 5 p.p.m.; the fragment mass tolerance was 20 p.p.m.; the oxidation of methionine was set as a variable modification and carbamidomethylation of cysteine as a fixed modification.

### Sample preparation and analysis of de-N-glycosylated EPO

EPO was denatured, reduced, alkylated and digested by PNGase F as mentioned above. De-N-glycosylated EPO samples were injected into a LC system (Waters ACQUITY, Miford, MA, USA), coupled with a hybrid quadrupole-orbitrap mass spectrometer. Solvent A was 0.1% FA and solvent B was 0.1% FA in ACN. Using a RP monolithic column (ProSwift™ RP-4H, 1×250 mm, Thermo Fisher Scientific, Bremen, Germany), solvent B started with 5% for a duration of 2 min, increasing to 25% in 7 min and to 60% in 34 min at a flow rate of 0.2 mL/min. Detailed MS instrument parameters are described in Supporting Information.

To deconvolute the MS raw data, the parameters of BioPharma Finder™ were set as follows: the amino acid sequence of epoetin beta was used as a FASTA database; the oxidation of methionine and carbamidomethylation of cysteine residues were included as variable modifications; the deamidation on Asn-24, −38 and −83 was set as a static modification. For the processing method, m/z range was set from 1,500 to 3,000; the Xtract (Isotopically Resolved) was used as a deconvolution algorithm; output mass range was set from 16,000 to 30,000; charge states between 8 and 25 were considered and a minimum number of 6 was used for deconvolution; the mass tolerance was set to 20 p.p.m..

## RESULTS AND DISCUSSION

### The identification and relative quantification of permethylated N-glycans

Various monosaccharides served as building blocks in the Python script to match the highly accurate MS1 data (Figure 1a). Furthermore, phosphorylation of glycans has been proved to play crucial roles in protein transport and human diseases, for example, phosphorylation of Man is involved to control the extracellular levels of leukemia inhibitory factor (LIF).^31^ However, phosphorylated glycans remain poorly characterized by MS platform. In the designed Python script (Figure S2), phosphorylated N-glycans were covered, which showed an increase of 93.98198 Da on Man (P-Hex) after permethylation (Figure 1b). Firstly, the A2F N-glycan standards were used to test this strategy, in which 52 N-glycan compositions (72 N-glycan structures) were identified with the characterization at MS1 and MS2 levels after permethylation (Table S1). In addition, 140 N-glycan compositions were identified from EPO-3 and EPO-4 samples, including 237 N-glycan structures in total (Table S2). Of them, different isomeric structures derived from one N-glycan composition were identified based on distinguishable retention time (RT) and MS2 fragments. For example, Neu5Ac_1_HexNAc_7_Hex_6_Fuc_1_Red-HexNAc_1_, extracted from N-glycans of EPO, showed six isomeric structures using C_18_ separation (Figure 1c). Trimannosylchitobiose core-free N-glycan species from EPO, containing a Red-HexNAc, were also considered for masses below 1,164.6251 Da that is the mass of trimannosylchitobiose core. They were also confidently identified at MS1 and MS2 levels although most species have no chitobiose cores for PNGase F cleavage (Figure S3).^32^ Eight phosphorylated N-glycans were identified from EPO samples, showing much higher coverage than the previous study.^33^ In this MS2 spectrum of HexNAc_1_Hex_5_P-Hex_1_Red-HexNAc_1_ (Figure 1d), the diagnostic fragment ion at m/z 313.1720 was generated from terminal P-Hex residue of the N-glycan by collision-induced dissociation (CID) fragmentation. However, N-glycans with O-acetylated sialic acids were not considered in Python scripts due to the removal of O-acetylation during permethylation. Taken together, the Python scripts were able to identify more N-glycan species compared to previous attempts.^24^

**Figure 1.**
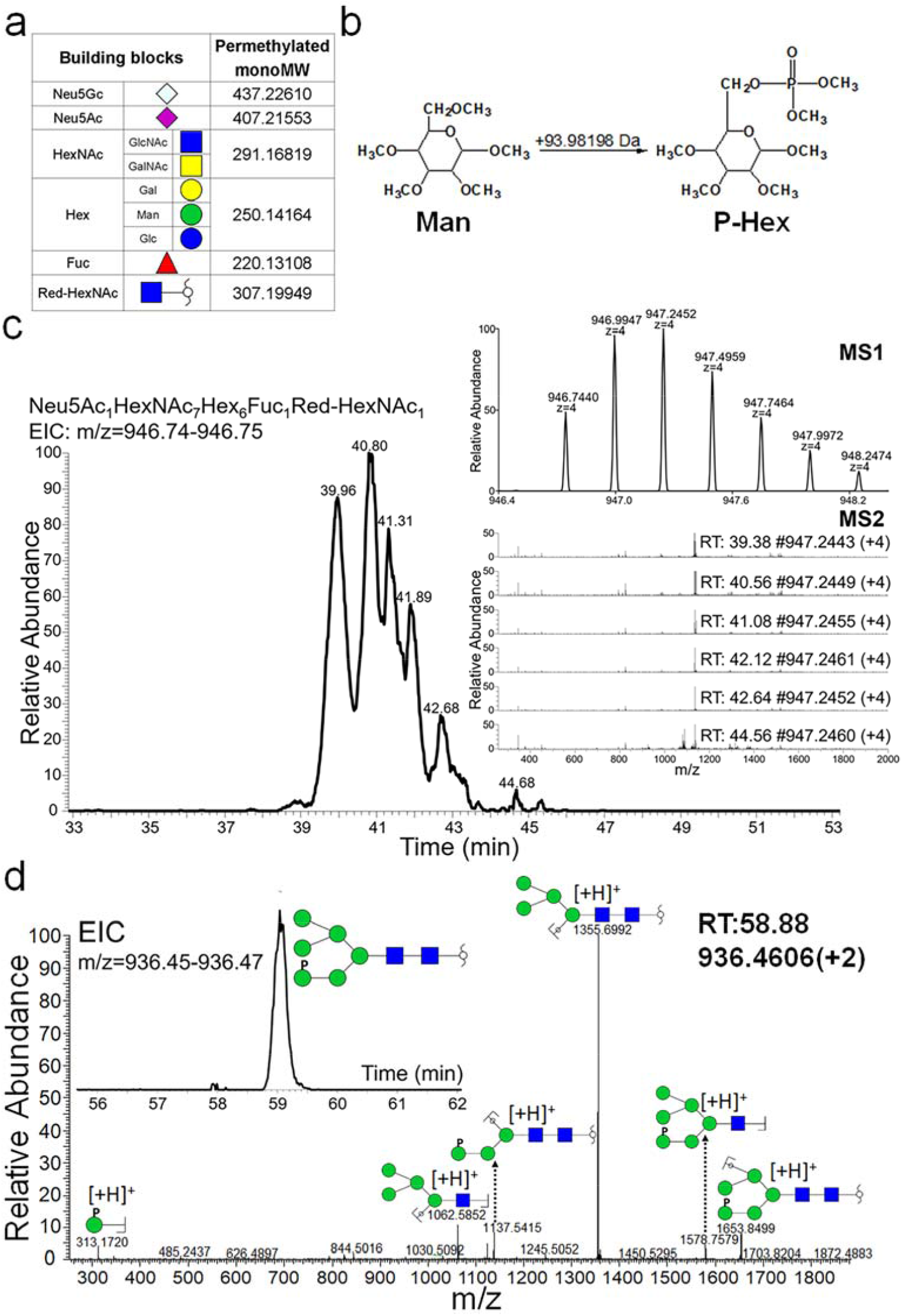
MS-based characterization of N-glycans with the newly designed Python scripts. (a) The basic building blocks used in the matching algorithm, showing each monoisotopic molecular weight (monoMW) after permethylation (GlcNAc, N-acetylglucosamine; GalNAc, N-acetylgalactosamine; Gal, galactose; Man, mannose; Glc, glucose). (b) The structures of permethylated P-Hex, as an additional building block compared to previous study. (c) The isomeric N-glycans separation by nanoLC(C_18_) and characterization using MS2 fragments, illustrated with the extracted Neu5Ac_1_HexNAc_7_Hex_6_Fuc_1_Red-HexNAc_1_ from the N-glycans of EPO. (d) The characterization of HexNAc_1_Hex_5_P-Hex_1_Red-HexNAc_1_ with EIC of precursors and the generated MS2 spectrum.

All N-glycan species were classified based on different modifications including sialylation and phosphorylation (Figure 2a). Most N-glycan species, 53%, were sialylated by Neu5Ac, 25% by both of Neu5Ac and Neu5Gc and 4% by Neu5Gc. The level of Neu5Gc contaminated N-glycans, 29% in total, is a key characteristic to evaluate the safety of the biotherapeutics for patient treatment, due to the immunogenic responses. In addition, 6% of all the identified N-glycan species were phosphorylated. To compare the amount of each N-glycan species, extracted ion chromatograms (EICs) of different charged species were integrated using Skyline and log2 transformed. All quantified N-glycan species of EPO-3 were summarized in Figure 2b, showing three highest abundant N-glycan species, Neu5Ac_4_HexNAc_5_Hex_7_Fuc_1_Red-HexNAc_1_, Neu5Gc_2_Neu5Ac_3_HexNAc_4_Hex_6_Fuc_1_Red-HexNAc_1_ and Neu5Ac_3_HexNAc_5_Hex_7_Fuc_1_Red-HexNAc_1_, and three lowest abundant N-glycan species, HexNAc_1_Hex_3_P-Hex_1_Red-HexNAc_1_, Neu5Gc_1_HexNAc_3_Hex_5_Red-HexNAc_1_ and HexNAc_2_Hex_3_Red-HexNAc_1_.

**Figure 2.**
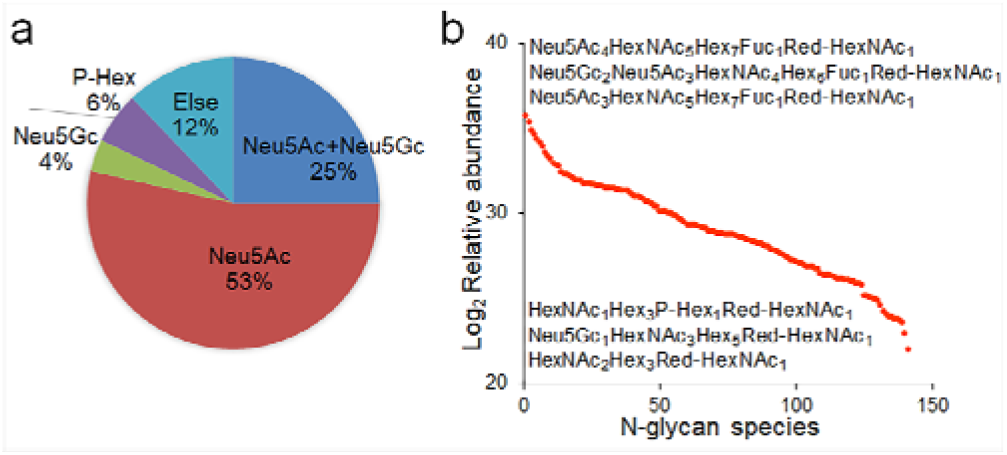
The classification and quantification analysis of identified N-glycan compositions of EPO. (a) The category analysis of identified N-glycan compositions of EPO based on the different modifications (sialylation and phosphorylation). (b) Comparative quantification of all the identified N-glycan compositions of EPO-3, showing the three highest (top) and three lowest (bottom) abundant N-glycan species.

### Batch-to-batch comparison of EPO derived N-glycans

In the production of therapeutic glycoproteins, the reproducibility of structural characteristics, such as PTMs, is highly important as it guarantees the continuous maintenance of pharmacological characteristics. Therefore, the N-glycans of two batches of EPO, EPO-3 and EPO-4, were compared to investigate the differential abundance of different N-glycan species across batches, as an essential step for quality control (QC). Using N-glycan MS data, total ion chromatograms (TICs) of both batches were firstly compared, showing a high similarity (Figure 3a). It has been proposed that Neu5Gc modifications of glycoprotein pharmaceuticals can lead to immunogenic responses of patients.^8^ During the biosynthesis of Neu5Gc, cytidine 5’-monophosphate (CMP)-Neu5Ac in the cytosol is the pre-cursor to generate CMP-Neu5Gc, catalyzed by CMP-Neu5Ac hydroxylase (CMAH).^34^ CMP-Neu5Gc is transferred into Golgi and then used to assemble glycoconjugates by various sialyltransferases. The ratio between Neu5Ac and Neu5Gc, as an index to quantify the generated Neu5Gc-containing N-glycans of EPO, was measured based on the peak areas of the MS2 diagnostic fragments, m/z 344.1704 for Neu5Ac and 374.1809 for Neu5Gc. The analysis revealed that the amount of Neu5Ac was about 18-fold larger compared to Neu5Gc in EPO-3 and 17-fold in EPO-4 (Figure 3b). After two-sample student’s T-test analysis by Perseus software (EPO-3 vs EPO-4), the significantly differential abundant N-glycan species were visualized in a volcano plot. The N-glycans, showing at least 1.5 between the two batches, were considered (Figure 3c). Neu5Ac_1_HexNAc_7_Hex_8_Fuc_1_Red-HexNAc_1_ (Green dot) showed a 1.7-FC higher abundancy in EPO-4 compared to EPO-3. In contrast, eleven N-glycan species showed at least 1.5-FC higher abundancy in EPO-3 compared to EPO-4 (1.5-FC < Blue dots < 2-FC and Red dots > 2-FC). To further investigate the N-glycans with abundance difference higher than 2-FC, six N-glycan species were compared in the heatmap (Figure 3d). As the most significant differences, HexNAc_2_Hex_3_Red-HexNAc_1_ was 4.85-FC higher in EPO-3 compared to EPO-4. Besides, most N-glycans, 128 species (91.4%), fluctuated within 1.5-FC between two batches of EPO samples, showing high stability and repeatability during EPO production.

**Figure 3.**
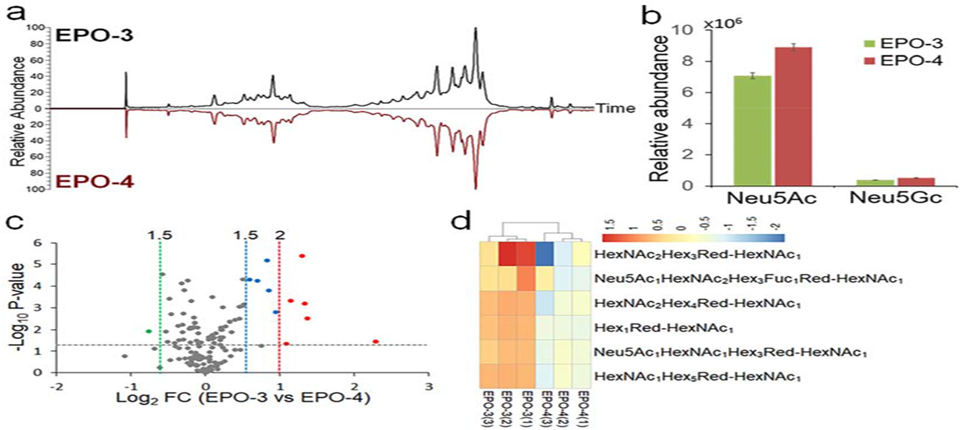
The quantitative comparison of identified N-glycan compositions between two different batches, EPO-3 and EPO-4. (a) The TIC comparison of N-glycans from EPO-3 and EPO-4. (b) The quantitative comparison of sialylation levels with Neu5Ac and Neu5Gc between EPO-3 and EPO-4. (c) The quantitative comparison of all identified N-glycan compositions by volcano plot (EPO-3 vs EPO-4). (d) The heatmap of significantly different abundance of identified N-glycan compositions with more than 2-FC (EPO-3 vs EPO-4).

### Peptide identification by pFind

Trypsin cleaves peptides at the C-terminus of Arg and Lys with a high site-specificity, except for -Arg/Lys-Pro-bonds. Chymotrypsin is less specific and cleaves peptide bonds at the C-terminus of Tyr, Phe, Trp and Leu. In addition, Met, Ala, Asp and Glu can be cleaved by chymotrypsin at a lower cleavage rate. Through the combination of chymotryptic and tryptic peptide analysis of EPO-3, a total sequence coverage of 74.1% was identified, using the pFind algorithm (Figure S4), in which 62% of the protein sequence was covered by tryptic peptides and 54.8% was covered by chymotryptic peptides. Cys-7 and −161 were modified with carbamidomethylation due to alkylation and Met-54 was identified with oxidation. Except for N-glycosylated peptides, most chymotryptic and tryptic peptides of EPO, were identified from MS data using the bottom-up approach. Also, non-O-glycosylated peptides, EAISPPDAAS(126)AAPLR from tryptic digestion and GAQKEAISPPDAAS(126)AAPL (Figure S5) from chymotryptic digestion, were identified. The analysis revealed that non-O-glycosylated EPO species existed and thus enabled the identification of O-glycans based on the mass difference between the non-O-glycosylated and O-glycosylated EPO species after de-N-glycosylation. Furthermore, the identified modifications were further considered in the deconvolution process of LC-MS data of de-N-glycosylated EPO.

### Glycopeptide analysis by pGlyco and the identification of O-acetylated sialic acids

In addition to the comprehensive qualitative and quantitative analysis of N-glycans, the characterization of glycopeptides enables the site-specificity of N-glycans on the protein backbone. The glycopeptide analysis was performed in technical triplicates using nanoLC(RP)-MS/MS and glycopeptides identified in at least two replicates were considered as valid data. After tryptic digestion of EPO-3, several EAEN(24)ITTGCcAEHCcSLNEN(38)ITVPDTK peptides was identified with one N-glycan species attached on Asn-24 or −38, while most of them, attached with two N-glycans, were not identifiable by pGlyco (Table S3). In this case, chymotryptic digestion showed a better applicability for glycopeptide identification compared to tryptic digestion as chymotryptic cleavage separated the Asn-24 and −38 on two different peptides (LL)EAKEAEN(24)ITTGCcAEHCcSL and NEN(38)ITVPDTKVNF(Y), which can be identified using pGlyco (Table S4). For EPO-3, the identified N-glycan compositions from chymotryptic glycopeptide and N-glycan analysis were compared in Table S5. On (LL)EAKEAEN(24)ITTGCcAEHCcSL peptides, 17 N-glycan compositions were identified by pGlyco, of whom 12 were also identified by permethylated N-glycan analysis. Parallelly, 13 N-glycan compositions were identified on NEN(38)ITVPDTKVNF(Y) peptides and 10 of them were identified by permethylated N-glycan analysis; 49 N-glycan compositions were identified on the (L)VN(83)SSQPW(EPL) peptides and 27 of them were confirmed by permethylated N-glycan analysis.

The O-acetylation of sialic acids was removed during permethylation for N-glycan analysis, while glycopeptide enabled the identification of O-acetylated sialic acids due to gentler sample preparation. However, pGlyco cannot identify these glycan modifications, which will promote next, necessarily required improvement of this software. For this reason, glycopeptides attached with O-acetylated sialic acid residues were able to be searched from MS data, with a mass increase of 42.01057 Da compared to non-O-acetylated species. As an example, the glycopeptide LVN(83)SSQPW(Neu5Ac_2_HexNAc_6_Hex_7_Fuc_1_), was identified by using pGlyco at m/z 1338.5282 (Figure 4a). With an addition of one O-acetyl group, LVN(83)SSQPW(Neu5Ac+OAc_1_Neu5Ac_1_HexNAc_6_Hex_7_F uc_1_) was identified with m/z 1352.5311 at MS1 level. Based on the diagnostic fragments, at m/z 316.1012, 334.1119 and 699.2452 could be identified at the MS2 level, evidencing the existence of Neu5Ac+OAc contaminated N-glycan species on the glycopeptide (Figure 4b).

**Figure 4.**
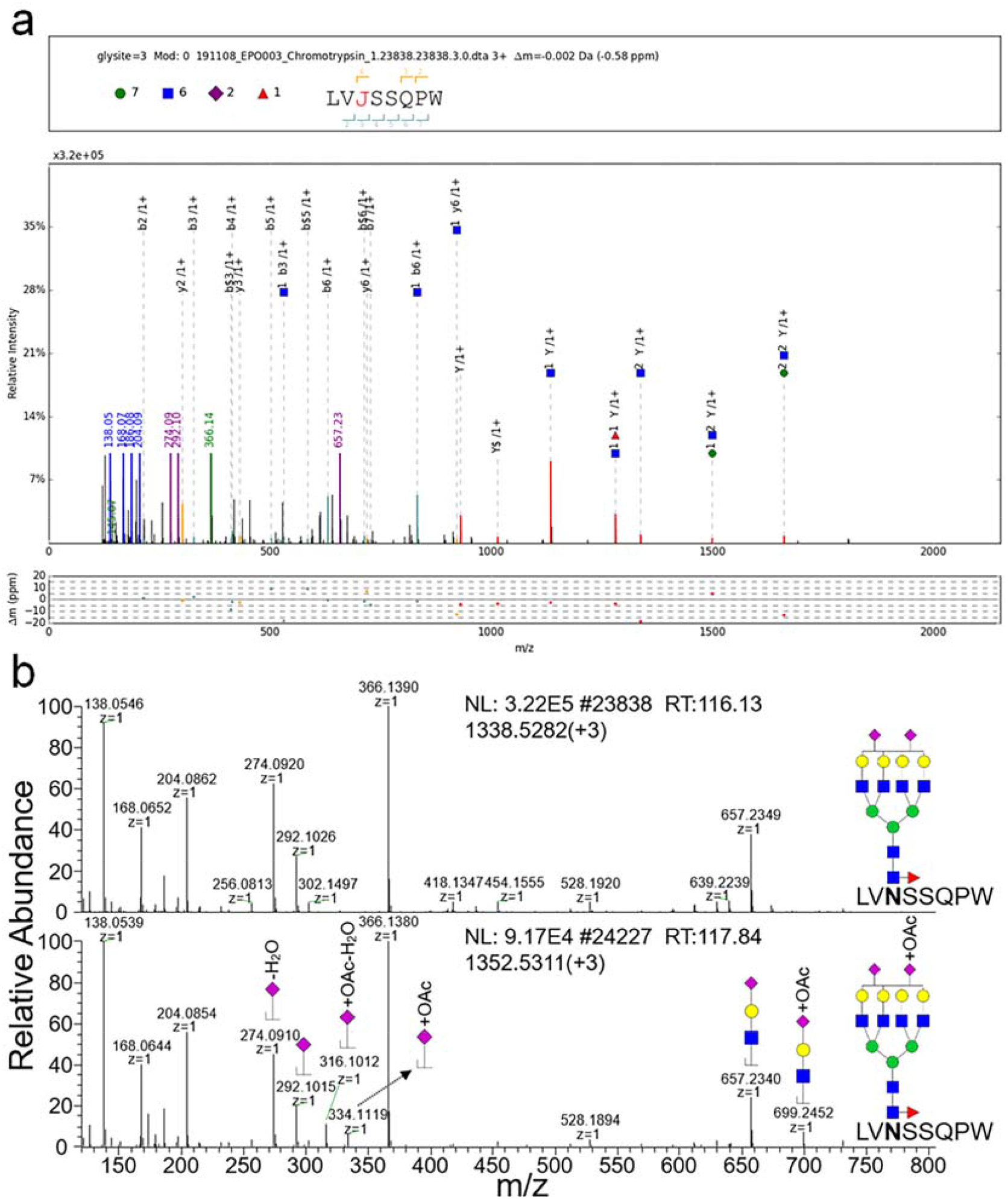
The identification of O-acetylated sialic acid by glycopeptide analysis. (a) The MS2 spectrum annotation of LVN(83)SSQPW(Neu5Ac_2_HexNAc_6_Hex_7_Fuc_1_), identified by pGlyco. (b) The comparison of LVN(83)SSQPW(Neu5Ac_2_HexNAc_6_Hex_7_Fuc_1_) and LVN(83)SSQPW(Neu5Ac+OAc_1_Neu5Ac_1_HexNAc_6_Hex_7_Fuc_1_) at MS1 and MS2 levels.

### The comparison of identified N-glycans between three approaches

This Python script-based analytical approach enabled the identification of 140 N-glycan compositions of EPO after OSPP preparation using MS data in this study. Using EPO-3, Chymotrypsin-based bottom-up analysis of glycopeptides identified 54 N-glycan compositions and a trypsin-based bottom-up approach identified 103 N-glycan compositions by pGlyco analysis. 29 N-glycan compositions were identified by all approaches; 51 species were exclusively identified by the analysis of tryptic glycopeptides, 10 by chymotryptic glycopeptide analysis and 101 by the analysis of permethylated N-glycans (Figure S6). The 29 N-glycan compositions were more abundant than other species and the small overlap between different approaches was mainly resulted from the low abundant glycopeptides or N-glycans, which was limited by switching from MS1 to MS2 level using data dependent acquisition (DDA). Using above three approaches, 215 N-glycan compositions were identified from EPO-3 in total.

### O-glycan identification by LC-MS analysis of de-N-glycosylated EPO

O-glycans are mostly released by chemical approaches such as β-elimination.^35^ As a limitation, these techniques can result in the degradation of the released O-glycans, significantly reducing the reliability of identified O-glycan species. For this reason, the investigation of O-glycans at the O-glycopeptide and O-glycoprotein level is advantageous, as it saves the intact O-glycans. Compared to N-glycans, mainly separated in three major types: high-mannose, complex and hybrid species with a trimannosylchitobiose core structure (HexNAc_2_Hex_3_),^36^ O-glycan shows a higher heterogeneity without a common core structure and lower site-specificity.^37^

However, O-glycans contribute much less to the heterogeneity of EPO with one O-glycosylation site (Ser-126) compared to N-glycans. LC-MS analysis of de-N-glycosylated EPO-3 revealed three main species with charges from 8 to 22 (Figure 5a). After deconvolution by BioPharma Finder™, three monoisotopic masses were identified. These includes 18459.584, 19115.795 and 19406.902 Da, and 18459.584 Da was identified as the monoisotopic mass of non-O-glycosylated EPO-3. The mass difference of 656.211 Da, between 18459.584 and 19115.795 Da species, revealed the O-glycan composition of Neu5Ac_1_HexNAc_1_Hex_1_ and a mass difference of 947.318 Da, between 18459.584 and 19406.902 Da species, revealed an O-glycan composition of Neu5Ac_2_HexNAc_1_Hex_1_ (Figure 5b).

**Figure 5.**
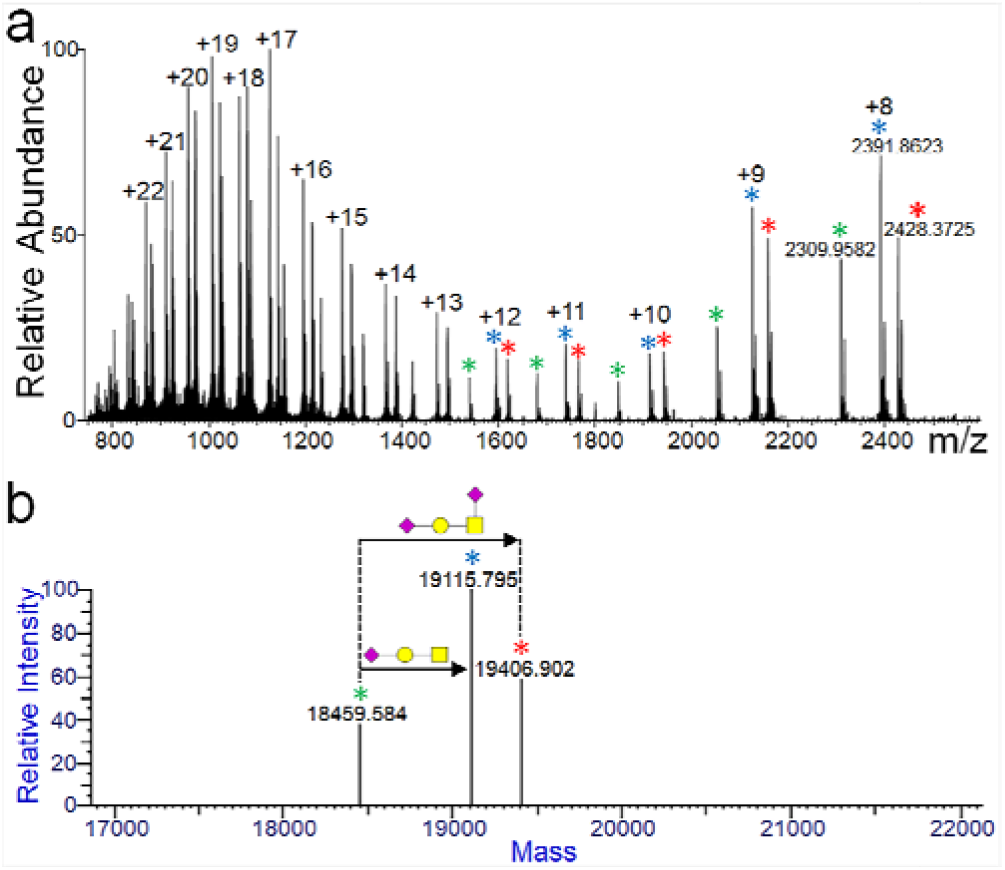
The analysis of de-N-glycosylated EPO-3 with three main species, labelled with green, blue and red asterisks. (a) MS1 spectrum of de-N-glycosylated EPO-3, showing three main species with charges from 8 to 22. (b) The detected monoisotopic masses after deconvolution of LC-MS data generated from de-N-glycosylated EPO-3 by BioPharma Finder™, showing the two major O-glycosylated species.

## CONCLUSION

In conclusion, an integrated strategy for in-depth glycosylation analysis, including permethylated N-glycan top-down analysis, glycopeptide bottom-up analysis using different proteases and O-glycan identification by LC-MS analysis of de-N-glycosylated proteins, was developed. For N-glycan analysis, the Python scripts were newly designed to allow the identification of the low abundant N-glycan species in commercial standard. Including phosphorylated Man residues, the additionally designed Python script enables the identification of phosphorylated N-glycans, showing powerful practicability in N-glycomics. The uncommon N-glycan species identified in this study, like trimannosylchitobiose core-free N-glycans, still need further investigation. Understanding different glycan patterns was significant to improve the QC procedure for the glycosylation of EPO, providing feedbacks to manage the EPO preparation, especially regarding the medically critical sialylation (Neu5Gc) level. For such detailed analysis, pGlyco alone is limited, as it cannot identify glycan modifications, like O-acetylation of sialic acids, and glycopeptides attached with multiple glycans. De-N-glycosylation reduced the heterogeneity of EPO species significantly and enabled the identification of the major O-glycan species.

The combination of these techniques and improvements in the newly developed strategy allows an in-depth characterization of glycan species from EPO. That can also be further applied to analyze the other therapeutic glycoproteins, with the aim to understand drug functions and side effects and improve the QC procedures.

## Supporting information

Preprint_Reference_24

## ASSOCIATED CONTENT

### Supporting Information

The Supporting Information is available free of charge on the ACS Publications website.

Supplementary Method Section, Tables (Tables S1, S2, S5) and Figures (Figure S1-S6). (PDF)

Supplementary Tables (Table S3 and S4). (XLSX)

## AUTHOR INFORMATION

### Notes

The authors declare no competing financial interest.

## ACKNOWLEDGMENT

Yudong Guan and Min Zhang were supported by the program of the China Scholarship Council (CSC No. 201606220045 and 201806780022).

## For Table of Contents Only

**Figure 1.**
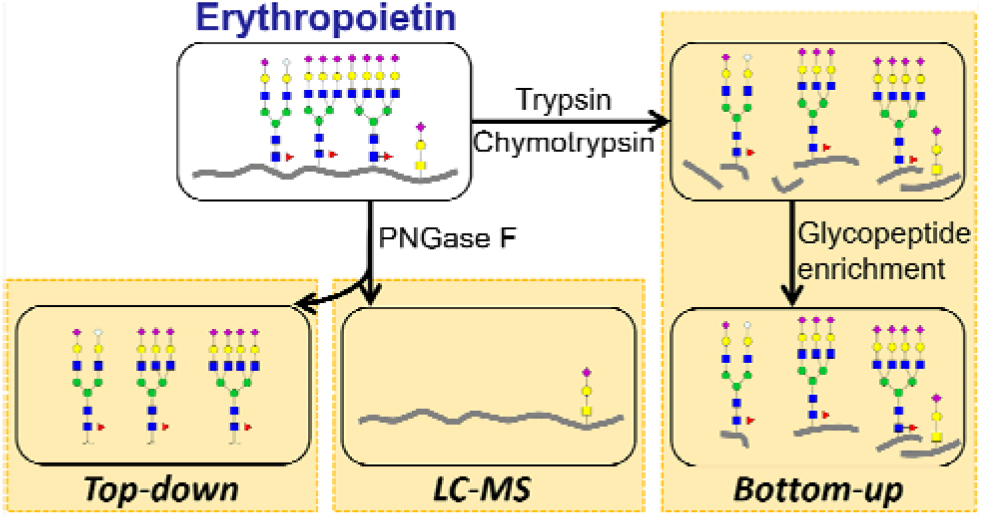

## Notes

### Competing Interest Statement

The authors have declared no competing interest.

