## Supplementary material for "An integrated strategy reveals complex glycosylation of erythropoietin using top-down and bottom-up mass spectrometry": Preprint_Reference_24

bioRxiv preprint doi: https://doi.org/10.1101/2020.10.15.340638; this version posted October 15, 2020. The copyright holder for this preprint (which was not certified by peer review) is the author/funder. All rights reserved. No reuse allowed without permission.

### An improved comprehensive strategy for deep and quantitative N-glycomics based on optimization of sample preparation, isotope-based data quality control and quantification, new N-glycan libraries and new algorithms

Yudong Guan[1], Jiaxiang Hu[2], Weiqian Cao[3], Wencong Cui[4], Fan Yang[5], Christoph Krisp[1], Ling Lin[3], Min Zhang[1], Hannah Voss[1], Raphael Schuster[6], Guoquan Yan[3], Marceline Manka Fuh[1], Morten Thaysen-Andersen[7], Nicolle H. Packer[7], Huali Shen[3], Pengyuan Yang[3] and Hartmut Schlüter[1*]

[1]Institute of Clinical Chemistry and Laboratory Medicine, University Medical Center Hamburg-Eppendorf, Hamburg, Germany.

[2]State Key Laboratory for Agrobiotechnology, China Agricultural University, Beijing, China.

[3]Institutes of Biomedical Sciences and Department of Chemistry, Fudan University, Shanghai, China.

[4]Department of Pediatric Hematology and Oncology, University Medical Center Hamburg-Eppendorf, Hamburg, Germany.

[5]Albrecht-Kossel-Institute for Neuroregeneration, University Medical Center Rostock, Rostock, Germany.

[6]Institute of Organic Chemistry, University of Hamburg, Hamburg, Germany.

[7]Department of Molecular Sciences, Macquarie University, Sydney, Australia.

*

bioRxiv preprint doi: https://doi.org/10.1101/2020.10.15.340638; this version posted October 15, 2020. The copyright holder for this preprint (which was not certified by peer review) is the author/funder. All rights reserved. No reuse allowed without permission.


#### ABSTRACT

Global in-depth analysis of N-glycosylation, as the most complex post-translational modification of proteins, is requiring methods being as sensitive, selective and reliable as possible. Here, an enhanced strategy for N-glycomics is presented comprising optimized sample preparation yielding enhanced glycoprotein recovery and permethylation efficiency, isotopic labelling for data quality control and relative quantification, integration of new N-glycan libraries (human and mouse), newly developed R-scripts matching experimental MS1 data to theoretical N-glycan compositions and bundled sequencing algorithms for MS2-based structural identification to ultimately enhance the coverage and accuracy of N-glycans. With this strategy the numbers of identified N-glycans are more than doubled compared with previous studies, exemplified by etanercept (more than 3-fold) and chicken ovalbumin (more than 2-fold) at nanogram level. The power of this strategy and applicability to biological samples is further demonstrated by comparative N-glycomics of human acute promyelocytic leukemia cells before and after treatment with all-trans retinoic acid, showing that N-glycan biosynthesis is slowed down and 57 species are significantly altered in response to the treatment. This improved analytical platform enables deep and accurate N-glycomics for glycobiological research and biomarker discovery.

bioRxiv preprint doi: https://doi.org/10.1101/2020.10.15.340638; this version posted October 15, 2020. The copyright holder for this preprint (which was not certified by peer review) is the author/funder. All rights reserved. No reuse allowed without permission.

#### INTRODUCTION

Glycosylated proteins, prevalent in all eukaryotic species[1], are involved in many biological processes such as recognition, signaling and communication between biomolecules and cells[2]. Altered protein glycosylation arising from physiology-dependent glycoprotein biosynthesis is associated with aberrant cellular and tissue physiology and may thus have diagnostic and prognostic value by being a potential reporter of disease[3-6]. Methods for getting views into the N-glycome as deep as possible are needed for a better understanding of protein glycosylation on a system-wide level.

Profiling of N- and O-glycans by mass spectrometry (MS) is yielding maps of the glycome, and demands relatively laborious specialized sample handling, data acquisition and informatics strategies. For example, the use of porous graphitized carbon (PGC)-liquid chromatography (LC)-MS/MS in negative polarity ion mode requires post-column make-up solvent supplement and manual annotation to achieve high glycome coverage[7]. A shortcoming of PGC-LC-MS/MS based profiling is the lack of a multiplexing option to perform comparative glycomics of multiple glycan samples within a single run and the limited analytical "transparency" of mono-, di- and tri-saccharides that do not retain well on analytical PGC columns. Other different analytical routes have been developed and used in glycomics, involving chemical derivatization such as permethylation[8]. Glycan permethylation is an often used glycan derivatization strategy due to several recognized benefits including: 1.) the stabilization of the glycan structures to prevent sample handling- and MS-induced decomposition, 2.) enhancement of their hydrophobicity for easier desalting, chromatographic separation on standard

bioRxiv preprint doi: https://doi.org/10.1101/2020.10.15.340638; this version posted October 15, 2020. The copyright holder for this preprint (which was not certified by peer review) is the author/funder. All rights reserved. No reuse allowed without permission.

reversed-phase LC columns and better MS properties including ionization in positive polarity mode, and 3.) an option for multiplexing enabling in-sample quantitative comparison of multiple glycan samples via isotopic labelling. Building on classical approaches for glycan permethylation, Mechref et al. developed a method for solid-phase permethylation of glycans in micro-columns packed with sodium hydroxide beads[9]. Known shortcomings of glycan permethylation include a degree of undesired side-reactions, incomplete derivatization, loss of modifications and partial glycan decomposition with a concomitant loss of sensitivity[10-12].

Glycoinformatics has seen exciting developments over the last two decades[13]. However, the bioinformatic solutions available for glycomics still lag behind the mature tools available for genomics and proteomics. Thus, glycoinformatics requires further improvements to be keep up with the analytical developments in glycomics and to be compatible with other "omics" related informatics solutions.

Aiming to enable a more efficient method for deep and quantitative N-glycomics, we first developed an optimized N-glycan preparation protocol (Supplementary Chapter 1), including optimized solid-phase permethylation (OSPP), second a novel informatics workflow (Supplementary Chapter 2), which employs R-script-based algorithms matching experimental high-accuracy MS1 data of permethylated N-glycans, extracted by MaxQuant[14], to extended human and mouse N-glycan libraries with isotope-based data quality control (Supplementary Chapter 3) and bundled sequencing algorithm for N-glycan structural characterization using MS2 data (Supplementary Chapter 4). Also, this designed workflow enables to perform isotopic labelling based

bioRxiv preprint doi: https://doi.org/10.1101/2020.10.15.340638; this version posted October 15, 2020. The copyright holder for this preprint (which was not certified by peer review) is the author/funder. All rights reserved. No reuse allowed without permission.

quantification (Supplementary Chapter 5) and relative quantification using Skyline and Perseus[15,16]. This novel N-glycomics strategy, which was orthogonally compared with identified glycan compositions at the glycopeptide level by pGlyco, shows a significant larger coverage and identification accuracy than existing glycan profiling methods as demonstrated for several isolated glycoproteins and complex mixtures of glycoproteins including protein extracts from human acute promyelocytic leukemia (APL) cells and mouse corpus callosum.

#### RESULTS

##### The optimized workflow for deep N-glycomics

Fig. 1 summarizes the optimized workflow for N-glycan profiling developed in this study and highlights the improvements at the levels of sample preparation and data analysis. We first optimized the initial sample preparation, including the (glyco)protein enrichment (Supplementary Chapter 1.1), N-glycan purification (Supplementary Chapter 1.2) and solid-phase permethylation based on existing techniques (Supplementary Chapter 1.3) (Supplementary Fig. 1-7) (Supplementary Table 1)[9,17-25], and then tested this optimized preparation approach using different glycoprotein and glycoproteome samples (Supplementary Chapter 1.4), showing simpler and more quantitative N-glycomics than 4-aminobenzoic acid butyl ester (ABBE)-based reductive amination (Supplementary Chapter 1.5) (Supplementary Fig. 8). We also improved the analytical strategy by using a new informatics workflow employing MS1 and MS2 data input (Fig. 1). MaxQuant in the version 1.6.2.3 (http://www.maxquant.org) was used to extract masses of all tentative glycan precursors in an “allPeptides.txt” file from the MS raw data, derived from $^{12}$C-

bioRxiv preprint doi: https://doi.org/10.1101/2020.10.15.340638; this version posted October 15, 2020. The copyright holder for this preprint (which was not certified by peer review) is the author/funder. All rights reserved. No reuse allowed without permission.

and $^{13}$C-permethylated N-glycans, respectively[26]. In this "allPeptides.txt" file, all the masses from the "Mass" column are extracted as "Mass.csv" and matched to the possible monosaccharide compositions using newly designed R-scripts, outputting the result file named as "Monosaccharide composition.csv" (Supplementary Chapter 2). In this glycoinformatics strategy, the data quality is controlled with stable isotopes, $^{12}$C and $^{13}$C, which serve to ensure a low false discovery rate (FDR) at MS1 level (Supplementary Chapter 3). To further realize structural N-glycomics after pairing between $^{12}$C- and $^{13}$C-permethylated species, bundled sequencing algorithm is developed to illustrate the N-glycan structure with the MS2 data (Supplementary Chapter 4).

The R-scripts consider tailored monosaccharide compositions for human and mouse N-glycans (https://github.com/guan181992/Glyco-informatics). For humans, the monoisotopic molecular weights (monoMWs) of $^{12}$C- and $^{13}$C-permethylated N-acetylneuraminic acid (Neu5Ac), N-acetylhexosamine (HexNAc), hexose (Hex), fucose (Fuc) and reduced N-acetylhexosamine (Red-HexNAc) are used as the building blocks to calculate the theoretical masses of N-glycans (Supplementary Fig. 9a). The deviation threshold of matching algorithm in R-script is firstly investigated based on a theoretical library of N-glycans below 6,000 Da containing the $HexNAc_1Hex_3Red\text{-}HexNAc_1$ (trimannosylchitobiose core) (Supplementary Table 2) (Supplementary Fig. 9b). The deviations of most species (10,380 from 10,618 species, 97.8%) are above 1.86 p.p.m. (Supplementary Fig. 10a) and 1.5 p.p.m. is used as the deviation threshold in the designed R-script in this study for $^{12}$C-permethylated N-glycan identification (Supplementary Fig. 11), in which multiple masses from MaxQuant can be matched with one targeted monosaccharide composition within ±1.5 p.p.m.

bioRxiv preprint doi: https://doi.org/10.1101/2020.10.15.340638; this version posted October 15, 2020. The copyright holder for this preprint (which was not certified by peer review) is the author/funder. All rights reserved. No reuse allowed without permission.

(Supplementary Fig. 10b). Also, the $^{13}C$-based library of N-glycans below 6,000 Da containing the $HexNAc_1Hex_3$Red-$HexNAc_1$ is matched to the experimental MS1 data and most of the deviations (9,950 from 9,995 species, 99.5%) show a mass deviation above 1.12 p.p.m. (Supplementary Table 3) (Supplementary Fig. 10c). Therefore, 1.1 p.p.m. is used as mass deviation threshold in the designed R-script for $^{13}C$-permethylated N-glycan identification. To cater for the MS1 data with different accuracies, the deviation thresholds of matching experimental masses in $^{12}C$- and $^{13}C$-based R-scripts are tested successfully from 1 to 5 p.p.m., showing more false positives with higher deviation thresholds. Furthermore, an additional R-script was designed to identify the N-glycans in mass range below 1,150 Da without considering trimannosylchitobiose core structure (Supplementary Fig. 11)

For analysis of mouse derived N-glycans, N-glycolylneuraminic acid (Neu5Gc) (Supplementary Fig. 10d), not synthesized in humans[27], is added as a building block when designing the N-glycan library. The mouse derived $^{12}C$-permethylated N-glycan library, containing the $HexNAc_1Hex_3$Red-$HexNAc_1$ up to 6,000 Da, totals 34,404 different monosaccharide compositions, in which 20,127 species (58.5%) have the deviation below 0.01 p.p.m. (Supplementary Table 4) (Supplementary Fig. 10e). $^{13}C$-permethylated N-glycan library also has similar dataset due to the isobaric $Neu5Ac_1Hex_1$ and $Neu5Gc_1Fuc_1$. Here, 1.5 p.p.m. is used as a mass tolerance in the $^{12}C$-based R-script and 1.1 p.p.m. in the $^{13}C$-based R-script with the addition of Neu5Gc, for mouse derived N-glycome identification, in which different monosaccharide compositions can be deduced from one precursor at MS1 level but allowed to be distinguished at MS2 level.

bioRxiv preprint doi: https://doi.org/10.1101/2020.10.15.340638; this version posted October 15, 2020. The copyright holder for this preprint (which was not certified by peer review) is the author/funder. All rights reserved. No reuse allowed without permission.

Pairs of $^{12}C$-/$^{13}C$-labelled species at the MS1 level has been applied on glycopeptide identification, enabling a low FDR[28]. In the MS raw data of 1:1 mixtures from permethylated N-glycans with $^{12}CH_3I$ and $^{13}CH_3I$, true and false positives were distinguished by pairs of retention time and peak area (roughly 1:1) between $^{12}C$- and $^{13}C$-permethylated species, exemplified by Enbrel-H (Supplementary Fig 12a, b, c). Using the "Monosaccharide composition.csv" generated by R-scripts, the FDR analysis was performed using the number of false positives divided by the number of all the matched precursors[29], in which Enbrel-H achieved an average of 62.5% for $^{12}C$-permethylated N-glycans from three replicates (Supplementary Table 6), 44.7% for $^{13}C$-permethylated N-glycans and 8.6% for their intersection of identification. Chicken ovalbumin, another model N-glycoprotein, achieved 60.1%, 42.4% and 10.3% respectively (Supplementary Fig. 12d). Therefore, the pairing of $^{12}C$/$^{13}C$ species reduced the N-glycan FDR significantly. In addition, with the monoMWs of building blocks, matching monoisotopic m/z were found to be another efficient strategy to improve the data quality.

For the monosaccharide compositions paired with $^{12}C$/$^{13}C$-labelled species at the MS1 level, MS2 fragments are extracted to characterize N-glycan structures using GlycoWorkbench in the version 2.1 stable build 146 (https://download.cnet.com/GlycoWorkbench-64-bit/3000-2383_4-75758804.html)[30]. At the MS2 level, the common MS2 diagnostic ions are summarized (Supplementary Table 5) and a novel algorithm of bundled sequencing is developed to enable fast structural characterization of N-glycans (Supplementary Fig. 13). The diagnostic ions, mainly fragmented as B- and Y-ions from defined bundled groups of different monosaccharides,

bioRxiv preprint doi: https://doi.org/10.1101/2020.10.15.340638; this version posted October 15, 2020. The copyright holder for this preprint (which was not certified by peer review) is the author/funder. All rights reserved. No reuse allowed without permission.

simplify the N-glycan identification significantly. In addition, the combination of diagnostic ions and bundled sequencing algorithm will promote advanced glycoinformatics development at MS2 level ( Supplementary Chapter 4). The N-glycan fragment annotations of the MS2 spectra follow the nomenclature proposed by Domon et al.[31].

**Application and validation of the analytical strategy**

Etanercept derived N-glycans have previously been characterized, however, these previous studies mainly focused on the highly abundant N-glycans[32,33]. In this study, each N-glycan sample from Enbrel-H and chicken ovalbumin were analyzed by both MALDI-MS and nanoLC-MS/MS, combining their specific advantages for glycan analysis. For Enbrel-H, 32 monosaccharide compositions were identified by MALDI-MS, by searching the permethylated N-glycan species with sodium adducts from the library mentioned above (Fig. 2a), while 90 monosaccharide compositions (162 N-glycan structures) were identified by nanoLC-MS/MS using our optimized analytical workflow (Supplementary Table 7). Without chromatographic separation, most N-glycans in low abundance were not identifiable due to suppressed ionization by MALDI-MS analysis. All the N-glycans identified by MALDI-MS were also present in the glycome profile obtained using nanoLC-MS/MS (Fig. 2b). The number of identified monosaccharide compositions was at least three-fold greater than other recent studies (Fig. 2c) and covered all the previously reported N-glycan species. For chicken ovalbumin, 23 monosaccharide compositions were identified by MALDI-MS (Fig. 2d), while 57 monosaccharide compositions (133 N-glycan structures) were identified by

bioRxiv preprint doi: https://doi.org/10.1101/2020.10.15.340638; this version posted October 15, 2020. The copyright holder for this preprint (which was not certified by peer review) is the author/funder. All rights reserved. No reuse allowed without permission.

nanoLC-MS/MS (Fig. 2e) (Supplementary Table 8), two-fold greater glycan coverage relative to other recent studies (Fig. 2f). With different chromatographic behaviors, isomeric species of the permethylated N-glycans were separated and identified by nanoRP($C_{18}$)-LC-MS/MS, which were unable to be distinguished by MALDI-MS. For example, Neu5$Ac_1$HexN$Ac_3$$Hex_4$$Fuc_1$Red-HexN$Ac_1$ from Enbrel-H was measured at m/z 2,417.229 [$M+Na^+$] by MALDI-MS (Supplementary Fig. 14a), while four isomers were separated by nanoRP($C_{18}$)-LC (Supplementary Fig. 14b, c), showing that each monosaccharide composition may derive to multiple N-glycan structures.

The permethylated N-glycome samples prepared in three replicates, APL-H from dimethylsulfoxide (DMSO)-treated APL cells and APL-6-H from all-trans retinoic acid (ATRA)-treated APL cells after 6 days (APL-6 cells), were also analyzed by nanoLC-MS/MS. Using this analytical strategy, 245 monosaccharide compositions (398 N-glycan structures) were identified from both cell groups (Supplementary Table 9). All N-glycans were characterizated at the MS1 and MS2 levels, almost showing the integral biosynthesis process in endoplasmic reticulum (ER) and Golgi apparatus (Supplementary Fig. 15).

pGlyco is an efficient software to analyze glycopeptides by reporting site-specific monosaccharide compositions. This tool was utilized in this study to verify the monosaccharide compositions identified in our newly designed workflow. For Enbrel-H, 33 monosaccharide compositions were identified by both approaches while 3 monosaccharide compositions, HexN$Ac_2$$Fuc_1$, HexN$Ac_1$$Fuc_1$ and HexN$Ac_3$$Hex_2$$Fuc_1$, were only identified at glycopeptide level by pGlyco (Fig. 3a). 57 species mostly at low abundance were exclusively identified at the N-glycan level due to the improvement of

bioRxiv preprint doi: https://doi.org/10.1101/2020.10.15.340638; this version posted October 15, 2020. The copyright holder for this preprint (which was not certified by peer review) is the author/funder. All rights reserved. No reuse allowed without permission.

ionization efficiency after OSPP preparation. The N-glycome of APL cells was also compared between OSPP-based N-glycan analysis and pGlyco-based glycopeptide analysis (Fig. 3b). 238 monosaccharide compositions were identified by pGlyco (Supplementary Table 10). 127 monosaccharide compositions were identified by both approaches; 111 species were exclusively identified by pGlyco, including the trimannosylchitobiose-free glycans that were beyond the consideration of R-sripts we developed, and 118 by OSPP-based glycan analysis. However, the isomeric N-glycans cannot be distinguished by pGlyco. The above data were analyzed by three triplicates and the identifications by more than once were considered valid.

To further test the power of this analytical strategy, the N-glycans from corpus callosum of an adult mouse were also analyzed. In total, 343 monosaccharide compositions (832 N-glycan structures) were identified (Supplementary Table 11). Of these, 64% were exclusively sialylated by Neu5Ac, 5% by Neu5Gc and 3% by both types of sialylation (Supplementary Fig. 16a). Quantitative analysis based on MS2 diagnostic ions of m/z 344.1704 (Neu5Ac) and 374.1809 (Neu5Gc) supported this finding by demonstrating that the amount of Neu5Gc was significantly lower (14%) than the level of Neu5Ac (normalized to 100%) (Supplementary Fig. 16b).

#### Quantitative and statistical analysis for differential N-glycomics

The differential N-glycomics was firstly performed by isotope-labelling quantification using MALDI-MS after OSPP preparation, with the N-glycome of APL cells permethylated by $CD_3I$ (APL-D) and $^{13}CH_3I$ (APL-13C). APL-D and APL-13C were compared with APL-6-H, respectively (Supplementary Chapter 5) (Supplementary Fig.

bioRxiv preprint doi: https://doi.org/10.1101/2020.10.15.340638; this version posted October 15, 2020. The copyright holder for this preprint (which was not certified by peer review) is the author/funder. All rights reserved. No reuse allowed without permission.

17, 18). In total, 56 monosaccharide compositions were quantified by MALDI-MS analysis (Supplementary Fig. 19). However, most N-glycans detected at low abundance could not be quantitatively profiled using this detection method due to their low signal strength and the low resolution of MALDI-MS.

Fig. 4 describes the workflow for relative quantification of the N-glycome we developed in this study. Relative quantification based on extracted ion chromatogram (EIC) data of LC-MS is tested for quantitative N-glycomics between APL-6-H and APL-H. The list of tentatively identified N-glycans is used as input into the Skyline software in the version 19.1 (http://skyline.maccosslab.org). The peak area of each monosaccharide composition (monoisotopic precursor m/z) is determined and used to establish the relative N-glycan abundance. The resulting output tables that are exported from Skyline included the monosaccharide composition and Total Area MS1 (the abundance of each monosaccharide composition) to input into Perseus in the version 1.6.2.1 (http://www.perseus-framework.org). Normalization and two-sample student's T-test were performed to compare the N-glycome quantification profile between APL and APL-6 cells, in which $p < 0.05$ and a fold change (FC) > 2 were defined as the minimum requirements to reach statistical significance.

After the treatment by DMSO (control group) and ATRA in DMSO solution on APL cells, quantitative N-glycomics and proteomics were performed, respectively (Fig. 5a). Between APL and APL-6 cells derived N-glycans, the biological replicates showed a high intra-group ($R^2 > 0.96$) and a low inter-group ($R^2 < 0.86$) by Pearson correlation analysis (Supplementary Fig. 20a). After two-sample student's T-test by Perseus (APL-6-H *vs* APL-H), the different abundance of N-glycans were visualized in a volcano

bioRxiv preprint doi: https://doi.org/10.1101/2020.10.15.340638; this version posted October 15, 2020. The copyright holder for this preprint (which was not certified by peer review) is the author/funder. All rights reserved. No reuse allowed without permission.

plot (Fig. 5b). In comparison with APL-H, the significantly up-regulated (red dots) and down-regulated (green dots) N-glycans of APL-6-H displaying a FC greater than 2 and $p < 0.05$ were identified. The 57 N-glycan species with significant differences in abundance were shown in a heat map by clustering analysis (Fig. 5c). Most significantly, the N-glycan precursors in APL-H were much higher than APL-6-H, including $HexNAc_1Hex_{8-11}Red\text{-}HexNAc_1$, showing that ATRA slowed down the N-glycan biosynthesis process. These quantitative differences were partly confirmed by the MALDI-MS analysis using isotope-based quantification (Supplementary Fig. 17, 18, see above). However, the Pearson correlation analysis based on the label-free quantification (LFQ) of proteome between APL and APL-6 cells showed an intra-group $R^2$ higher than 0.97 and inter-group $R^2$ lower than 0.80 (Supplementary Fig. 20b). During N-glycan biosynthesis, α-glucosidase II (α-Glc II), a heterodimeric enzyme including catalytic and regulatory subunits (UniProtKB accession number: Q14697 and P14314) and responsible for the generation of $HexNAc_1Hex_{9-10}Red\text{-}HexNAc_1$[44], was changed as about 1.1-fold in APL-6 cells compared with APL cells (Fig. 5d). This enzyme showed much less sensitivity than $HexNAc_1Hex_9Red\text{-}HexNAc_1$ and $HexNAc_1Hex_{10}Red\text{-}HexNAc_1$, changed as 21% and 12% in APL-6 cells compared to APL cells respectively, during ATRA medication of APL cells (Fig. 5e).

#### DISCUSSION

We have described an enhanced strategy for deep and quantitative N-glycomics, in which the N-glycan sample preparation is improved including the OSPP of the oligosaccharides, yielding highly purified N-glycans and simplified structures. Focusing on the

bioRxiv preprint doi: https://doi.org/10.1101/2020.10.15.340638; this version posted October 15, 2020. The copyright holder for this preprint (which was not certified by peer review) is the author/funder. All rights reserved. No reuse allowed without permission.

permethylated N-glycans, a glycoinformatics solution for in-depth analysis at MS1 and MS2 levels was developed. The narrow deviation threshold in the matching algorithm of R-script and isotopic-labelling enables high data quality for N-glycan identification at MS1 level. Significantly, the permethylated N-glycans increase the number of identifiable precursor masses, obtaining high coverage of the N-glycome with generation of simple fragments by collision-induced dissociation (CID) to allow the fast identification of N-glycan structures by bundled sequencing algorithm. In comparison to previous studies, this novel strategy was shown to identify a larger number of N-glycans because with the improved method N-glycans in the lower abundance range are detected, even with starting sample amounts at the nanogram level in the case of purified glycoproteins and at the microgram level in the case of complex protein mixture. The identified monosaccharide compositions were verified by pGlyco software at the glycopeptide level. Besides in-depth identification, global relative quantification for comparative N-glycomics was also demonstrated via nanoLC-MS/MS by integration of EICs, showing more advantages than stable isotopic labelling approach using MALDI-MS. With differential quantitative N-glycomics by nanoLC-MS/MS it was shown that the effect of the ATRA medication on N-glycan biosynthesis of APL cells is more obvious than the view on the changes of the proteins, partly due to the smaller dataset of N-glycome in the cells. The analytical approach described here has the potential to adapt it for in-depth structural O-glycomics. The demonstrated glycan profiling for human and mice samples can be extended to other mammalian species, with the N-glycan library developed in this study.

This improved strategy for deep and quantitative N-glycomics will be beneficial to

bioRxiv preprint doi: https://doi.org/10.1101/2020.10.15.340638; this version posted October 15, 2020. The copyright holder for this preprint (which was not certified by peer review) is the author/funder. All rights reserved. No reuse allowed without permission.

identify N-glycan biomarkers and improve our understanding of the structure, biosynthesis and function of the N-glycome.

#### ACKNOWLEDGMENTS

Y.G., W.C., and F.Y. were supported by the program of the Chinese Scholarship Council (CSC No. 201606220045, 201708080210, 201206170169). N.H.P., H.S., M.T.A. and P.Y. received funding support for the trilateral partnership (MQ-FU-HAM Trilateral Strategic Network) supported by a grant of the DAAD (Deutscher Akademischer Austausch Dienst, German Academic Exchange Service; Project ID: 57172988).

#### AUTHOR CONTRIBUTIONS

Y.G. and H.S. conceived and designed the study. Y.G. developed the algorithms, performed the experiments, analyzed the results and wrote the manuscript. J.Hu. contributed with the R-script writing. Y.G., W.C., L.L. and G.Y. performed the glycopeptide analysis. W.C. provided the APL cells and APL-6 cells. F.Y. provided the corpus callosum of an adult mouse. M.Z. and H.V. confirmed the high reproducibility of experiments and data analysis. R.S. contributed to the measurement of glycan by MALDI-MS. C.K., M.M.F., M.T.A., N.H.P., P.Y. and H.S. provided critical feedback for this study and manuscript.

#### COMPETING FINANCIAL INTERESTS

The authors declare no competing financial interests.

bioRxiv preprint doi: https://doi.org/10.1101/2020.10.15.340638; this version posted October 15, 2020. The copyright holder for this preprint (which was not certified by peer review) is the author/funder. All rights reserved. No reuse allowed without permission.

19. Zhou, S., Dong, X., Veillon, L., Huang, Y. & Mechref, Y. LC-MS/MS analysis of permethylated N-glycans facilitating isomeric characterization. *Anal. Bioanal. Chem.*

bioRxiv preprint doi: https://doi.org/10.1101/2020.10.15.340638; this version posted October 15, 2020. The copyright holder for this preprint (which was not certified by peer review) is the author/funder. All rights reserved. No reuse allowed without permission.

**409**, 453-466 (2017).

bioRxiv preprint doi: https://doi.org/10.1101/2020.10.15.340638; this version posted October 15, 2020. The copyright holder for this preprint (which was not certified by peer review) is the author/funder. All rights reserved. No reuse allowed without permission.

37. Szigeti, M. & Guttman, A. Automated N-glycosylation sequencing of

bioRxiv preprint doi: https://doi.org/10.1101/2020.10.15.340638; this version posted October 15, 2020. The copyright holder for this preprint (which was not certified by peer review) is the author/funder. All rights reserved. No reuse allowed without permission.

biopharmaceuticals by capillary electrophoresis. *Sci. Rep.* **7**, 11663 (2017).

bioRxiv preprint doi: https://doi.org/10.1101/2020.10.15.340638; this version posted October 15, 2020. The copyright holder for this preprint (which was not certified by peer review) is the author/funder. All rights reserved. No reuse allowed without permission.

#### Figure Legends

Figure 1. Overview about the newly designed workflow for in-depth N-glycomics and the areas in which optimizations were introduced, including the sample preparation and data analysis, to enable highly enriched N-glycans and the combination of software and algorithms for deep N-glycome profiling using MaxQuant, custom-made R-scripts, GlycoWorkbench and isotope-based data quality control based on high-resolution nanoLC-MS/MS precursor (MS1) and fragment data (MS2) of permethylated N-glycans.

Figure 2. Identified N-glycans derived from the etanercept, using Enbrel-H product in this study, and chicken ovalbumin by MALDI-MS and nanoLC-MS/MS. (a) Identified monosaccharide compositions of Enbrel-H by MALDI-MS after OSPP-based preparation (GlcNAc is abbreviated from N-Acetylglucosamine). (b) Veen diagram comparison of identified monosaccharide compositions between MALDI-MS and nanoLC-MS/MS analysis for Enbrel-H in this study. (c) Comparison of the numbers of identified monosaccharide compositions from Enbrel-H by nanoLC-MS/MS analysis after OSPP-based preparation in this study and other recent studies. (d) Identified monosaccharide compositions of chicken ovalbumin by MALDI-MS after OSPP-based preparation. (e) Veen diagram comparison of identified monosaccharide compositions between MALDI-MS and nanoLC-MS/MS analysis for chicken ovalbumin in this study. (f) Comparison of the numbers of identified monosaccharide compositions from chicken ovalbumin by nanoLC-MS/MS analysis after OSPP-based preparation in this study and other recent studies.

Figure 3. Comparative analysis of monosaccharide compositions between OSPP-based

bioRxiv preprint doi: https://doi.org/10.1101/2020.10.15.340638; this version posted October 15, 2020. The copyright holder for this preprint (which was not certified by peer review) is the author/funder. All rights reserved. No reuse allowed without permission.

N-glycomics and glycopeptide analysis with pGlyco software. (a) The comparison of identified monosaccharide compositions from Enbrel-H by OSPP-based N-glycan identification and pGlyco-based glycopeptide analysis. (b) The comparison of identified monosaccharide compositions from APL cells by OSPP-based N-glycan identification and pGlyco-based glycopeptide analysis.

Figure 4. Workflow of relative quantification of the N-glycome using Skyline and Perseus software.

Figure 5. Differential N-glycomics and proteomics of APL and APL-6 cells investigating the response of the cells towards ATRA treatment. (a) Overview of this experimental workflow for APL cells after DMSO and ATRA treatment respectively, showing that differential quantitative N-glycomics was performed by tribrid orbitrap mass spectrometer with CID fragmentation and differential quantitative proteomics was performed by hybrid orbitrap mass spectrometer with higher energy collisional dissociation (HCD) fragmentation. (b) Volcano plot of differential N-glycomics of APL-6-H *vs* APL-H. (c) Heat map of monosaccharide compositions present at significantly different abundance in APL-6-H *vs* APL-H. (d) Volcano plot of differential proteomics of APL-6 *vs* APL cells. (e) The quantitative comparison of differential N-glycan precursors and their related specific glucosidase, α-Glc II including Q14697 and P14314, in APL-6 *vs* APL cells.

bioRxiv preprint doi: https://doi.org/10.1101/2020.10.15.340638; this version posted October 15, 2020. The copyright holder for this preprint (which was not certified by peer review) is the author/funder. All rights reserved. No reuse allowed without permission.

Figure 1.

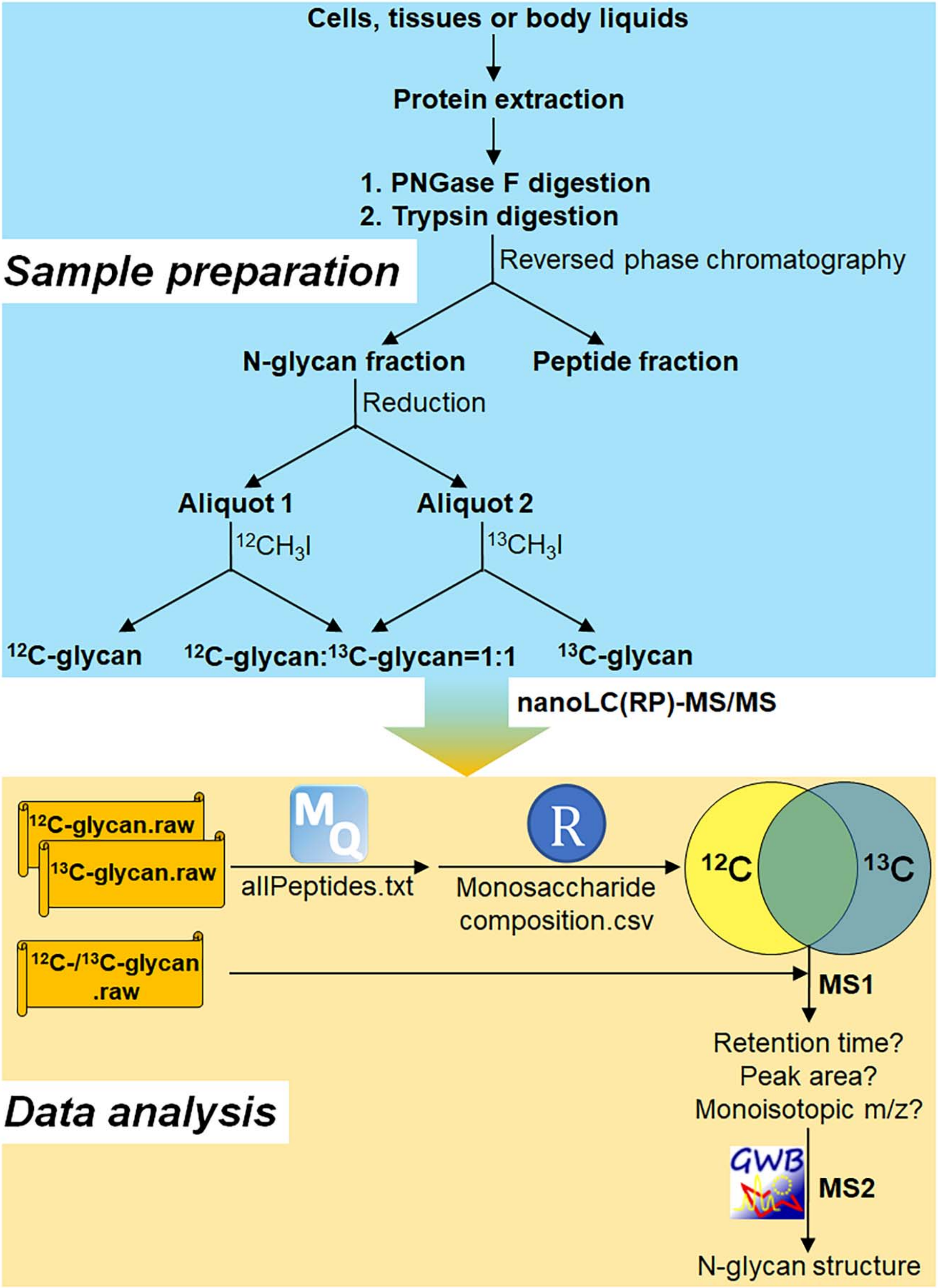

Figure 2.

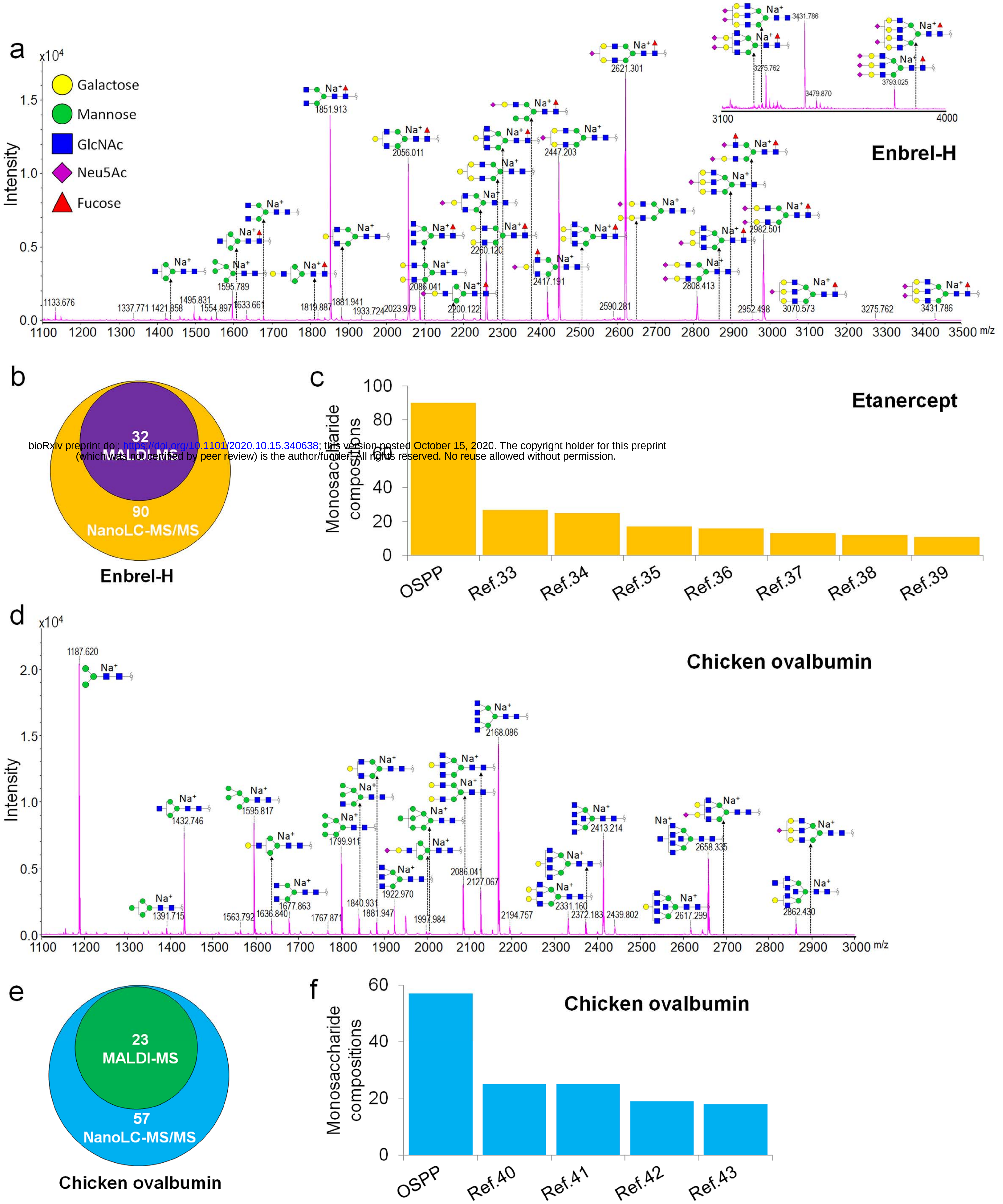


bioRxiv preprint doi: https://doi.org/10.1101/2020.10.15.340638; this version posted October 15, 2020. The copyright holder for this preprint (which was not certified by peer review) is the author/funder. All rights reserved. No reuse allowed without permission.

bioRxiv preprint doi: https://doi.org/10.1101/2020.10.15.340638; this version posted October 15, 2020. The copyright holder for this preprint (which was not certified by peer review) is the author/funder. All rights reserved. No reuse allowed without permission.

Figure 3.

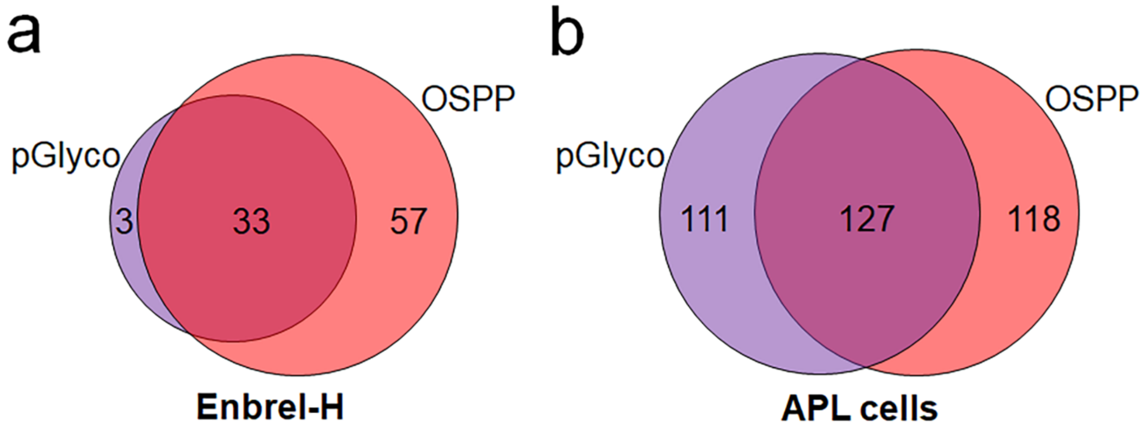

bioRxiv preprint doi: https://doi.org/10.1101/2020.10.15.340638; this version posted October 15, 2020. The copyright holder for this preprint (which was not certified by peer review) is the author/funder. All rights reserved. No reuse allowed without permission.

Figure 4.

**List of identified N-glycans**

| m/z | Mass | Neu5Ac | HexNAc | Hex | Fuc | Red-HexNAc | Deviation (p.p.m.) |
|---|---|---|---|---|---|---|---|
| 1,165.6331 | 1,164.6258 | 0 | 1 | 3 | 0 | 1 | 0.58 |
| 915.4914 | 1,828.9683 | 0 | 3 | 3 | 1 | 1 | 0.73 |
| .. | .. | .. | .. | .. | .. | .. | .. |
| .. | .. | .. | .. | .. | .. | .. | .. |
| 987.5118 | 2,959.5134 | 2 | 3 | 5 | 1 | 1 | 0.15 |
| .. | .. | .. | .. | .. | .. | .. | .. |

↓

| Monosaccharide composition | m/z | Charge | Mass |
|---|---|---|---|
| HexNAc$_1$Hex$_3$Red-HexNAc$_1$ | 1,165.6331 | 1 | 1,164.6258 |
| HexNAc$_3$Hex$_3$Fuc$_1$Red-HexNAc$_1$ | 915.4914 | 2 | 1,828.9683 |
| .. | .. | .. | .. |
| .. | .. | .. | .. |
| Neu5Ac$_2$HexNAc$_3$Hex$_5$Fuc$_1$Red-HexNAc$_1$ | 987.5118 | 3 | 2,959.5134 |
| .. | .. | .. | .. |

↓ Skyline (integration of EICs)

**List of N-glycans showing their abundance**

↓ Perseus (statistical analysis)

**List of N-glycans showing significant changes in their abundance**

Figure 5.

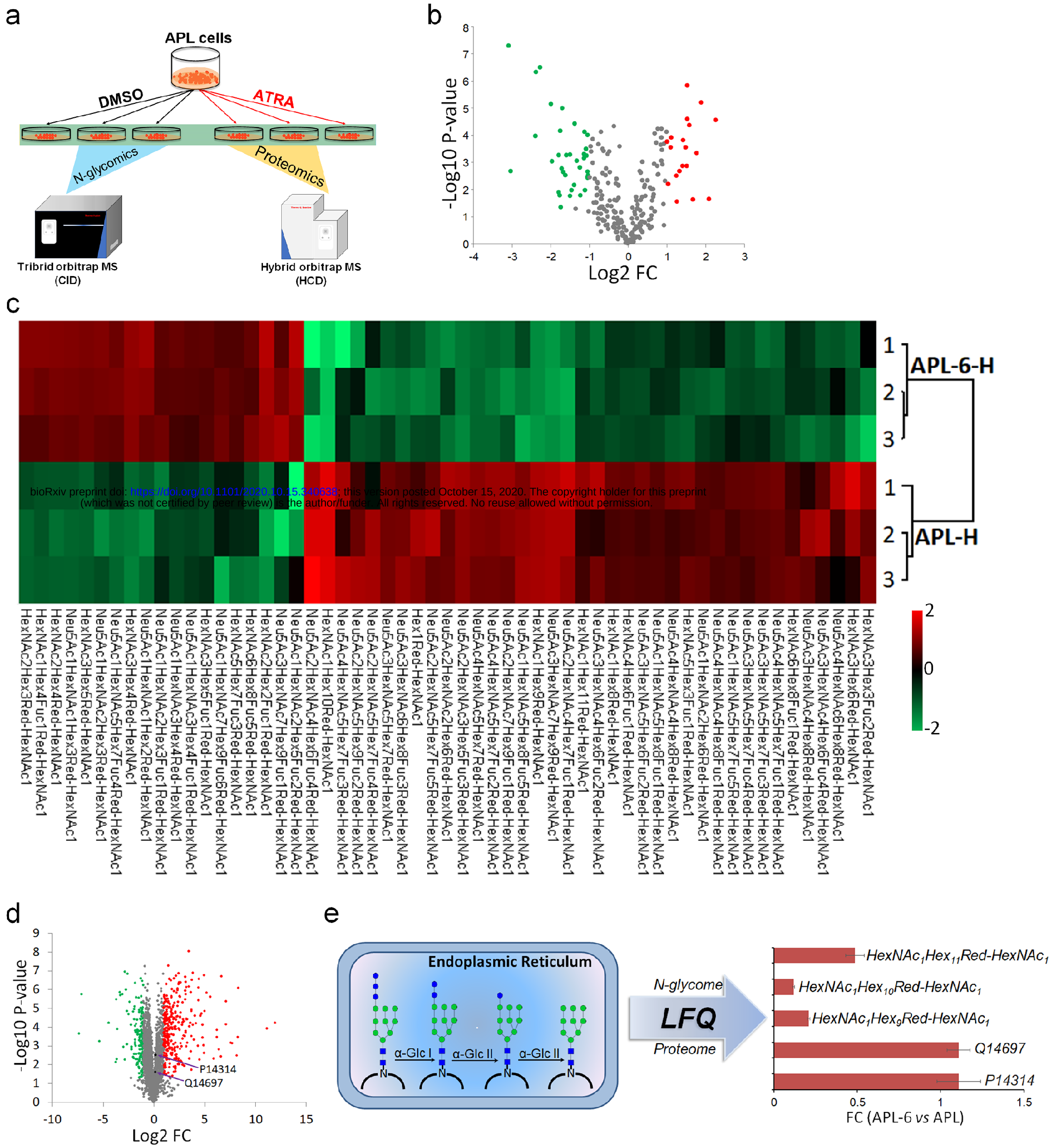


bioRxiv preprint doi: https://doi.org/10.1101/2020.10.15.340638; this version posted October 15, 2020. The copyright holder for this preprint (which was not certified by peer review) is the author/funder. All rights reserved. No reuse allowed without permission.

bioRxiv preprint doi: https://doi.org/10.1101/2020.10.15.340638; this version posted October 15, 2020. The copyright holder for this preprint (which was not certified by peer review) is the author/funder. All rights reserved. No reuse allowed without permission.

#### ONLINE METHODS

**Materials.** Phosphate-buffered saline (PBS) was purchased from Thermo Fisher Scientific (Bremen, Germany). Chicken ovalbumin, sodium hydroxide, dimethylsulfoxide (DMSO), iodomethane, iodomethane-$d_3$ ($CD_3I$), iodomethane-$^{13}C$ ($^{13}CH_3I$), sodium deoxycholate (SDC), tetraethylammonium bromide (TEAB) and all-trans retinoic acid (ATRA) were purchased from Sigma (Darmstadt, Germany). All other chemicals were also purchased from Sigma unless otherwise stated. Two different lots of Enbrel® (G30909, H17609) (etanercept) were purchased from Pfizer (Istanbul, Turkey), which were herein named Enbrel-G and Enbrel-H. Sequencing grade modified trypsin and PNGase F were obtained from Promega (Madison, WI, USA). 0.5 mL centrifugal filters (3 k and 10 k devices) were purchased from Merck KGaA (Darmstadt, Germany). Solid-phase-extraction (SPE) columns containing reversed-phase (RP) materials ($C_{18}$ Sep-Pak cartridges) were obtained from Waters (Miford, MA, USA). Mice (BALB/c) were purchased from the Charles River Laboratories (Wilmington, MA, USA). All experiments were approved by the local ethical committee and conducted according to the guidelines for the Care and Use of Laboratory Animals.

**Cell lines and cell culture.** Acute promyelocytic leukemia (APL) is distinguished from other forms of acute myeloid leukemia (AML) by its responsiveness to ATRA therapy. Treatment with ATRA allows DNA transcription and differentiation of immature leukemic promyelocytes into mature granulocytes by targeting the oncogenic transcription factor, promyelocytic leukemia-retinoic acid receptor α (PML-RARα). A NB4 APL-derived cell line (German Collection of Microorganisms and Cell Cultures,

bioRxiv preprint doi: https://doi.org/10.1101/2020.10.15.340638; this version posted October 15, 2020. The copyright holder for this preprint (which was not certified by peer review) is the author/funder. All rights reserved. No reuse allowed without permission.

Braunschweig, Germany) was cultured in RPMI 1640, supplemented with 10% FCS in a 95/5% air/$CO_2$ atmosphere. For neutrophil-like differentiation, cells were treated with 2 μM ATRA in DMSO for 6 days into three groups, hereafter referred to as "APL-6 cells", meanwhile, three control groups were treated with DMSO, hereafter referred to as "APL cells". The cells were washed twice using PBS solution before protein extraction.

**Protein extraction from cell lines and murine corpus callosum tissues.** 1 mL SDC buffer (1% SDC in 0.1M TEAB) was added to each cell sample followed by incubation at 99°C for 10 min and sonicated with 25% of normal energy at 3 cycles for 30 sec on ice. For corpus callosum homogenization, the isolated forebrain of an adult mouse was transected. The corpus callosum was dissected from adjacent tissues and frozen by liquid nitrogen immediately. Samples were homogenized in 150 μL 8 M urea and 50 μL SDC buffer and sonicated with 25% of normal energy at 3 cycles for 30 sec on ice[45]. The protein concentration was estimated using the Pierce™ BCA Protein Assay Kit, following the manufacturer's instructions (Thermo Fisher Scientific, Bremen, Germany).

**Tryptic digestion and desalting of the extracted proteins from APL cells.** 250 μg proteins, extracted from APL cells, were denatured in 6 M urea, reduced by 20 mM dithiothreitol (DTT) at 60°C for 30 min and alkylated by 40 mM iodoacetamide (IAA) at room temperature for 30 min in the dark. DTT and IAA were prepared in 100 mM ammonium bicarbonate (ABC) buffer. The samples were digested by trypsin (1:100, w/w) at 37°C for 20 h and quenched by adding 2% (v/v) formic acid (FA). Tryptic peptides were desalted using RP-SPE columns as described by Villén et al.[46]. The column was conditioned using 9 mL acetonitrile (ACN) and equilibrated with 5 mL 0.1% (v/v)

bioRxiv preprint doi: https://doi.org/10.1101/2020.10.15.340638; this version posted October 15, 2020. The copyright holder for this preprint (which was not certified by peer review) is the author/funder. All rights reserved. No reuse allowed without permission.

trifluoroacetic acid (TFA). Tryptic peptides were loaded onto the column and washing was performed, using 3 mL 0.1% (v/v) TFA. Finally, peptides were eluted by 3 mL 90% ACN in 0.5% (v/v) acetic acid. Eluted peptides were lyophilized and stored at -20° until further use.

**Different protein precipitation methods for protein recovery from APL cells.** Protein precipitation was performed in order to separate proteins from other biomolecule species, such as free glycans. Here, different precipitation methods were compared to the above protein precipitation-independent tryptic digestion (further referred to as SDC-based digestion), to estimate the highest protein extraction efficiency, prior to PNGase F-based N-glycan cleavage for glycome analysis.

250 μg proteins extracted from SDC-lysed APL cells were additionally precipitated using either chloroform/methanol or trichloroacetic acid (TCA) precipitation after alkylation[47,48]. For chloroform/methanol precipitation, four-fold volume of methanol was added to the alkylated protein solution. Afterwards, one-fold volume of chloroform and three-fold volume of water was added. After centrifugation for 10 min at 12,000 rpm using a Centrifuge 5424 (Eppendorf AG, Hamburg, Germany), the supernatant was discarded and three-fold volume of methanol was added. After an additional centrifugation step for 10 min at 12,000 rpm using a Centrifuge 5424, precipitated proteins were dried in a SpeedVac™ vacuum concentrator and stored at -20°C until further use.

TCA precipitation was performed additionally. The solution containing 250 μg proteins was mixed with an equivalent volume of 20% (v/v) TCA after alkylation. The mixture was incubated at -20°C for 1 h. Then samples were thawed at room temperature and the

bioRxiv preprint doi: https://doi.org/10.1101/2020.10.15.340638; this version posted October 15, 2020. The copyright holder for this preprint (which was not certified by peer review) is the author/funder. All rights reserved. No reuse allowed without permission.

supernatant was removed after centrifugation at 12,000 rpm by Centrifuge 5424 for 10 min at a temperature of 4°C. 0.5 mL ice-cold acetone was added and precipitated proteins were collected after centrifugation at 12,000 rpm by Centrifuge 5424 for 10 min at 4°C and dried in a SpeedVac™ vacuum concentrator. Dried proteins from methanol/chloroform and TCA precipitation were resuspended in 100 mM ABC buffer and digested by trypsin (1:100, w/w) at 37°C for 20 h. Resulting peptides were dried in a SpeedVac™ vacuum concentrator and stored at -20° until further use. All samples were analyzed using nanoLC-MS/MS and technical triplicates for each condition were measured.

**NanoLC-MS/MS analysis and raw data processing for differential proteomics.** Prior to LC-MS/MS analysis, tryptic peptides were dissolved in 0.1% (v/v) FA, transferred to an autosampler and injected into a Dionex Ultimate 3000 UPLC system (Thermo Fisher Scientific, Bremen, Germany). Peptides were purified and desalted using an RP $C_{18}$ trapping column (Thermo Scientific™ Acclaim PepMap™, 100 µm×2 cm, 5 µm, 100Å) at a flow rate of 15 µL/min with 1% solvent B (0.1% (v/v) FA in ACN) and 99% solvent A (0.1% (v/v) FA) and then transferred to an analytical RP $C_{18}$ column (Thermo Scientific™ Acclaim PepMap™ RSLC, 75 µm×50 cm, 2 µm, 100Å) after 10 min at a flow rate of 0.275 µL/min with 2% solvent B, for chromatographic separation. Peptides were separated through a linear gradient from 2 to 20% solvent B in 115.5min, from 20 to 32% in 135.5 min and from 32 to 95% in 136.5 min. Eluted peptides were ionized using a nano spray ion source for electrospray ionization at a capillary voltage of 1.8 kV. Peptide ions were transferred to a hybrid quadrupole-orbitrap mass spectrometer (Q Exactive, Thermo Fisher Scientific, Bremen, Germany). For MS1 scanning, the maximum injection

bioRxiv preprint doi: https://doi.org/10.1101/2020.10.15.340638; this version posted October 15, 2020. The copyright holder for this preprint (which was not certified by peer review) is the author/funder. All rights reserved. No reuse allowed without permission.

time was 60 ms for an AGC target of $3\times10^6$; m/z scan range was set from 375 to 1,600 with an orbitrap resolution of 70,000 FWHM at m/z 200 for data acquisition. Data dependent acquisition was performed in top N mode. For HCD-MS/MS, the 10 highest abundant precursor ions were selected for fragmentation with a normalized HCD collision energy of 27%; fragment spectra were recorded with the maximum injection time of 50 ms for an AGC target of $1\times10^5$ in an orbitrap mass analyzer with an orbitrap resolution of 17,500 FWHM at m/z 200.

Obtained nanoLC-MS/MS raw data were visualized using the Xcalibur 4.0.27.13 (Thermo Fisher Scientific, Bremen, Germany) and processed with Proteome Discoverer in the version 2.0 (Thermo Fisher Scientific, Bremen, Germany) using the SEQUEST algorithm. For peptide identification, obtained MS2 spectra were searched against theoretical fragment spectra of tryptic peptides, generated from the reviewed SWISSProt FASTA database, containing 20,239 entries, obtained in October 2018. For protein identification, the following parameters were used: the precursor mass tolerance was set to 10 p.p.m.; the fragment mass tolerance was set to 0.02 Da; variable modifications including oxidation (M, +15.995 Da) and acetyl (Protein N-term, +42.011 Da) were considered and the carbamidomethylation (C, +57.021 Da) was set as a fixed modification; minimum length of considered peptides was set to 6 amino acids; 2 missed tryptic cleavages were tolerated. A false discovery rate (FDR) value threshold <0.01, using a reverted decoy peptide databases approach, was set for peptide identification. For LFQ, the proteomics data from APL and APL-6 cell groups were processed by MaxQuant with the FASTA database in the version 1.6.2.3 (http://www.maxquant.org). The "unique plus razor peptides" was chosen for protein quantification. The precursor mass tolerance was set to 20 p.p.m.; the

bioRxiv preprint doi: https://doi.org/10.1101/2020.10.15.340638; this version posted October 15, 2020. The copyright holder for this preprint (which was not certified by peer review) is the author/funder. All rights reserved. No reuse allowed without permission.

fragment mass tolerance was 0.5 Da; dynamic modifications included oxidation, acetyl and fixed modification carbamidomethylation. Perseus software in the version 1.6.2.1 (http://www.perseus-framework.org) was utilized to visualize the data of differential proteins. Basically, the proteins labeled by only identified by site, reverse, and potential contaminant were removed from the data. Pearson correlations were calculated with $R^2$. Normalization and two-sample student's T-test were performed to compare the protein quantification profile between APL and APL-6 cells, in which FDR of 0.05 and a fold change (FC) > 2 were set.

**N-glycan release, purification and solid-phase permethylation.** 200 μg Enbrel-G (etanercept) was denatured, reduced and alkylated as described above. The buffer was exchanged to 100 mM ABC buffer using a prepacked 3 kDa stage tip filter. For glycan release, 1:30 (v/w) PNGase F was added and incubated at 37°C for 24 h. In order to optimize the N-glycan purification, the PNGase F digestion of Enbrel-G was divided into three aliquots, which were convicted to three different purification approaches, namely: 90% (v/v) ethanol precipitation, stage tip filter-based separation and tryptic digestion coupled with RP-SPE $C_{18}$ cartridge clean-up (Supplementary Fig. 1a). For 90% (v/v) ethanol precipitation, ABC buffer was firstly evaporated by a SpeedVac™ vacuum concentrator due to its effect on protein precipitation. De-N-glycosylated proteins were precipitated by adding 500 μL 90% (v/v) ethanol and removed by centrifugation at 12,000 rpm using Centrifuge 5424 for 10 min[19]. For filter separation, 10 kDa centrifuge filter tips were used to separate the N-glycans (filtrate fraction) from the larger de-N-glycosylated proteins (retentate fraction)[20]. For the tryptic digestion coupled with RP-SPE $C_{18}$ cartridge clean-up, the PNGase F digestion was first quenched by boiling at

bioRxiv preprint doi: https://doi.org/10.1101/2020.10.15.340638; this version posted October 15, 2020. The copyright holder for this preprint (which was not certified by peer review) is the author/funder. All rights reserved. No reuse allowed without permission.

99°C for 5 min. After cooling at room temperature, trypsin (1:100, w/w) was added and incubated at 37°C for 20 h. N-glycans and tryptic peptides were separated as described by Morelle et al. using a RP-SPE $C_{18}$ cartridge[21]. The RP cartridge was conditioned with 5 mL methanol and equilibrated with 10 mL 5% (v/v) acetic acid respectively. Then the digested sample was loaded into the cartridge and N-glycans were eluted with 5 mL 5% (v/v) acetic acid. The peptides were eluted by 90% ACN in 5% (v/v) acetic acid, stored in -20°C for subsequent O-glycan analysis. Purified N-glycan samples and an additional blank sample were permethylated using classical solid-phase permethylation as described by Mechref et al. (Supplementary Fig. 1b)[9]. Sodium hydroxide beads were used to pack spin columns, which were washed by 100 μL DMSO twice. Each purified N-glycan sample or the blank sample was redissolved in 50 μL DMSO with 1 μL water and 30 μL iodomethane. Samples were loaded into the spin column and re-loading was performed five times. Afterwards, 100 μL DMSO was loaded to wash the beads. 150 μL 5% (v/v) acetic acid were added into the collected fraction to quench the permethylation and eliminate oxidation reactions. Next, 200 μL chloroform was added and chloroform-water extraction was repeated ten times for each sample. Finally, each chloroform extraction was evaporated using a SpeedVac™ vacuum concentrator and stored at -20° until further use.

**The development of optimized solid-phase permethylation (OSPP).** The classical "solid-phase" permethylation approach describes permethylation based on sodium hydroxide powders or beads as a stationary phase inside a capillary or spin column. The approach is limited by complicated operation, limitation of water accessibility in the solid-phase sodium hydroxide and impracticality for multiple samples. In our

bioRxiv preprint doi: https://doi.org/10.1101/2020.10.15.340638; this version posted October 15, 2020. The copyright holder for this preprint (which was not certified by peer review) is the author/funder. All rights reserved. No reuse allowed without permission.

optimization, a solution containing water, DMSO and iodomethane was added to sodium hydroxide beads in a glass vial, to prevent the absorption of moisture from air during the rotation (Supplementary Fig. 1c). Briefly, the sodium hydroxide beads were weighed (200 mg) in a glass vial. Purified and dried N-glycans were redissolved in a water/DMSO solution and 100 µL iodomethane was added. Redissolved samples were transferred to the glass vial containing sodium hydroxide beads. Samples were incubated in a Thermomixer compact (Eppendorf AG, Hamburg, Germany) using a rotation speed of 1,300 rpm. The development of OSPP preparation was based on cleaved and purified N-glycans from Enbrel-G. Glycan permethylation was mainly performed by two different approaches: slurry and solid-phase permethylation. For slurry permethylation, a slurry of sodium hydroxide in DMSO was grounded in a dry mortar, added to released and dried glycans and mixed with 100 µL iodomethane[22]. For the assessment of the effects of different sodium hydroxide concentrations in DMSO, N-glycans cleaved from 20 µg Enbrel-G were dissolved with different sodium hydroxide concentrations (10, 20, 30, 60, 120 and 200 µg/µL) in 200 µL DMSO and 100 µL iodomethane was added (Experiment A). Each mixture was shaken for 10 min at 1,300 rpm using a Thermomixer compact and 200 µL 5% (v/v) acetic acid was added to quench the permethylation reaction. Permethylated glycans were extracted using 300 µL chloroform by chloroform-water extraction. For OSPP development, dried N-glycans cleaved from 20 µg Enbrel-G were dissolved in 110 µL water/DMSO (10/100, v/v) and 100 µL iodomethane was added into the mixture. Redissolved N-glycans were transferred to a glass vial containing 200 mg sodium hydroxide beads and shaken for 10 min at 1,300rpm by Thermomixer compact, which was compared to the slurry permethylation performed as described above, using

bioRxiv preprint doi: https://doi.org/10.1101/2020.10.15.340638; this version posted October 15, 2020. The copyright holder for this preprint (which was not certified by peer review) is the author/funder. All rights reserved. No reuse allowed without permission.

120 µg/µL of sodium hydroxide (Experiment B). OSPP was performed using different ratios of water/DMSO (0/100, 5/100, 10/100, 15/100, 20/100) (v/v), in the following referred to as Experiment C. Furthermore, different reaction times (10 min and 30 min) were compared in OSPP at a water/DMSO ratio of 10/100 (v/v) (Experiment D). Also, the addition of iodomethane before ($CH_3I$/NaOH) and after sodium hydroxide (NaOH/$CH_3I$) addition were compared (Experiment E). Finally, N-glycans were reduced by borane-ammonia to eliminate anomers before permethylation and compared to non-reducing N-glycan permethylation (Experiment F). Here, 10 µg/µL borane-ammonia were added to dried N-glycans and incubated at 60°C for 1 h[19]. Borane-ammonia was removed by evaporation with three additions of 300 µL methanol prior to permethylation (Compare Supplementary Table 1).

After above OSPP, the reaction solution was transferred into a new vial. 150 µL DMSO was added to further wash the sodium hydroxide beads and transferred into the new vial to extract permethylated N-glycans. 200 µL 5% (v/v) acetic acid were added to permethylated glycan solution to quench the reaction and eliminate oxidation reactions[10]. Then 300 µL chloroform were added and chloroform-water extraction was repeated ten times. Chloroform was evaporated using a SpeedVac™ vacuum concentrator. From the experiments described above, the optimal experimental parameters for OSPP were estimated and used for further experiments.

**4-aminobenzoic acid butyl ester (ABBE)-based reductive amination of N-glycans.** Reductive amination is another effective derivative method to improve glycan ionization efficiency for MS analysis, whose applicability was to be compared to glycan

bioRxiv preprint doi: https://doi.org/10.1101/2020.10.15.340638; this version posted October 15, 2020. The copyright holder for this preprint (which was not certified by peer review) is the author/funder. All rights reserved. No reuse allowed without permission.

permethylation by mass spectrometric analysis in this work. ABBE was used as a reductive amination reagent following the classical approach as described by Ruhaak et al.[49]. Briefly, 30 µL 0.35 M ABBE dissolved in acetic acid/DMSO (3/7, v/v) were added to dried N-glycans, cleaved from 20 µg Enbrel-G. 30 µL 1M 2-picoline borane prepared in DMSO was added and the reductive amination was incubated at 65°C for 1 h. The reaction was quenched by adding 900 µL water and evaporated using a SpeedVac™ vacuum concentrator. For comparison, equal amounts of N-glycans from Enbrel-G permethylated by OSPP and aminated with ABBE were injected into nanoLC-MS/MS as described below.

**MS analysis of permethylated and ABBE aminated N-glycans.** Prior to nanoLC-MS/MS analysis, permethylated and ABBE aminated N-glycans were dissolved into 0.1% (v/v) FA, transferred to an autosampler and injected into a Dionex Ultimate 3000 UPLC system (Thermo Fisher Scientific, Bremen, Germany). Derivative N-glycans were purified and desalted using an RP $C_{18}$ trapping column (Thermo Scientific™ Acclaim PepMap™, 100 µm×2 cm, 5 µm, 100Å) at a flow rate of 3 µL/min with 2% solvent B (0.1% (v/v) FA in ACN) and 98% solvent A (0.1% (v/v) FA) and transferred to an analytical RP $C_{18}$ column (Thermo Scientific™ Acclaim PepMap™ RSLC, 75 µm×50 cm, 2 µm, 100Å), at a flow rate of 0.2 µL/min, for chromatographic separation. For permethylated N-glycans, a 90 min gradient was used, starting with 10% solvent B. Solvent B increased to 30% in 5 min followed by a linear gradient elevating the concentration of 75% in 70 min and finally increased to 95% in 80 min. For ABBE aminated N-glycans, the gradient started with 5% solvent B for a duration of 10 min and then solvent B increased to 80% in 90 min.

bioRxiv preprint doi: https://doi.org/10.1101/2020.10.15.340638; this version posted October 15, 2020. The copyright holder for this preprint (which was not certified by peer review) is the author/funder. All rights reserved. No reuse allowed without permission.

Eluted, derivated N-glycans were ionized using a nano spray ion source for electrospray ionization at a capillary voltage of 1.8 kV. Derivative N-glycan ions were transferred to a tribrid quadrupole-orbitrap-ion trap mass spectrometer (Fusion, Thermo Fisher Scientific, Bremen, Germany). For MS1 scanning, an orbitrap mass analyzer was used with an orbitrap resolution of 120,000 FWHM at m/z 200; the maximum injection time was 120 ms, to an AGC target of $2\times10^5$; m/z scan range was from 450 to 2,000. Data dependent acquisition was used in the top speed mode. For CID-MS/MS, the most intense precursor ions were selected for fragmentation and isolated using an isolation window of 3; the normalized collision energy of CID was set to 35%; fragment ions were injected to an ion trap with maximum injection time was 20 ms at an AGC target of $1\times10^5$. The parallelization of MS1 and MS2 acquisition was performed as described by Senko et al.[50]. The data were visualized and analyzed using Xcalibur software.

**Matrix-assisted laser desorption ionization (MALDI)-MS analysis.** For MALDI-MS analysis, permethylated N-glycans were resuspended into 10 mM sodium chloride in 50% (v/v) methanol. Samples were dropped on a MALDI target with saturated α-cyano-4-hydroxycinnamic acid (CHCA) in 50% (v/v) methanol. The MALDI-TOF-TOF system (Ultraflextreme, Bruker, Bremen, Germany) was used for N-glycan analysis with a m/z range from 500 to 8,000. A MALDI laser energy was set to 30% in the positive-reflectron mode and analyzed using the flexAnalysis software in the version 3.3 (Build 80) (Bruker, Bremen, Germany).

**Glycopeptide enrichment and measurement by nanoLC-MS/MS.** Glycopeptides were enriched from 400 μg Enbrel-H, 1 mg proteins extracted from APL cells and APL-6 cells

bioRxiv preprint doi: https://doi.org/10.1101/2020.10.15.340638; this version posted October 15, 2020. The copyright holder for this preprint (which was not certified by peer review) is the author/funder. All rights reserved. No reuse allowed without permission.

digested by trypsin and resuspended in 100 μL 80% ACN with 1% (v/v) TFA. Glycopeptides were enriched as described by Liu et al.[30]. A micro-column was packed with 30 mg zwitterionic hydrophilic interaction liquid chromatography (ZIC-HILIC) particles obtained from a HILIC column (SeQuant® ZIC®-cHILIC 3μm, 100Å 250×4.6 mm, Merck KGaA, Darmstadt, Germany) on top of $C_8$ membrane materials (3M, Eagan, MN, USA). Column equilibration was performed with 600 μL 80% ACN with 1% (v/v) TFA. Samples were loaded and low-hydrophilic peptides were eliminated by adding 600 μL 80% ACN with 1% (v/v) TFA. Glycopeptides were eluted by 300 μL 0.1% (v/v) TFA and dried by a SpeedVac™ vacuum concentrator.

Lyophilized samples were redissolved in 0.1% (v/v) FA prior to nanoLC-MS/MS analysis in triplicates. Glycopeptides were transferred to an autosampler and injected into an EASY-nano-LC system (Thermo Fisher Scientific, Bremen, Germany) without the usage of a trap column. Samples were directly separated on an analytical RP $C_{18}$ column (Thermo Scientific™ Acclaim PepMap™ RSLC, 75 μm×50 cm, 2 μm, 100Å). Solvent A was 0.1% (v/v) FA and solvent B was 0.1% (v/v) FA in ACN. The gradient for Enbrel-H lasted for 1 h. The nano-pump started with 1% solvent B at a flow rate of 0.2 μL/min, increased to 20% in 40 min, reached 30% in 47 min and jumped at 90% in 50 min. For glycopeptides of APL and APL-6 cells, a 6 h gradient was used. The nano-pump started with 1% solvent B. Then, solvent B increased to 20% in 300 min, to 30% in 342 min and jumped to 90% in 345 min. Eluted glycopeptides were ionized using a nano spray ion source for electrospray ionization at a capillary voltage of 1.8 kV and were transferred to a tribrid quadrupole-orbitrap-ion trap mass spectrometer. The MS parameters were set as following: for MS1 scanning, the m/z scan range was set from 350 to 2,000; an orbitrap

bioRxiv preprint doi: https://doi.org/10.1101/2020.10.15.340638; this version posted October 15, 2020. The copyright holder for this preprint (which was not certified by peer review) is the author/funder. All rights reserved. No reuse allowed without permission.

mass analyzer was used with an orbitrap resolution of 120,000 FWHM at m/z 200; the maximum injection time was 50 ms, to a AGC target of $5\times10^5$; precursor ions with a charge state between 2 and 6 were considered. Data dependent acquisition was used in top speed mode. For HCD-MS/MS, the most intense precursor ions were selected for fragmentation and isolated, using an isolation window of 4; dynamic exclusion of selected precursor ions was performed for a duration of 15 s; isolated precursor ions were fragmented using a stepped HCD gradient with normalized collision energies of 20%, 30% and 40%; fragment ions were analyzed in an orbitrap mass analyzer at an orbitrap resolution of 15,000 FWHM at m/z 200; the maximum injection time was 250 ms, to an AGC target of $5\times10^5$.

Glycopeptides were identified from raw data by pGlyco in the version 2.1.2 (http://pfind.ict.ac.cn/software/pGlyco/index.html). The glycan database was extracted from GlycomeDB (www.glycome-db.org) with the total N-glycan entries of 7,884[30]. A protein database, containing 20,239 entries, was obtained from SWISSProt in October 2018. The amino acid sequence of etanercept was also utilized as the database FASTA file for its glycopeptide analysis. The parameters were set as following: the precursor tolerance was 5 p.p.m.; the fragment mass tolerance was 20 p.p.m.; as a variable modification, the oxidation of methionine was considered; carbamidomethylation of cysteine was set as a fixed modification.

bioRxiv preprint doi: https://doi.org/10.1101/2020.10.15.340638; this version posted October 15, 2020. The copyright holder for this preprint (which was not certified by peer review) is the author/funder. All rights reserved. No reuse allowed without permission.
