## Supporting Information for "An integrated strategy reveals complex glycosylation of erythropoietin using top-down and bottom-up mass spectrometry"

Supplementary Method Section

**MS instrument parameters for permethylated N-glycan analysis**

MS parameters were set as follows: nano spray ion source was set at a capillary voltage of 1.8 kV. for MS1 scanning, an orbitrap mass analyzer was used with a resolution of 120,000 full width at half maximum (FWHM) at m/z 200; the maximum injection time was 120 ms and AGC target was 2×10^5^; m/z scan range was set from 450 to 2,000. Data dependent acquisition (DDA) mode was performed in top speed mode. For collision-induced dissociation (CID)-MS/MS, the most intense precursor ions were selected for fragmentation and isolated with an isolation window of 3; the normalized collision energy of CID was set to 35%; fragment ions were injected to an ion trap with maximum injection time of 20 ms and AGC target was 1×10^5^.

**MS instrument parameters for peptide analysis**

MS parameters were set as follows: nano spray ion source was set at a capillary voltage of 1.8 kV. for MS1 scanning, the maximum injection time was 100 ms and AGC target was 5×10^5^; m/z scan range was set from 400 to 2,000 with an orbitrap resolution of 70,000 FWHM at m/z 200 for data acquisition. DDA was performed in top N mode. For higher-energy collisional dissociation (HCD)-MS/MS, the 15 highest abundant precursor ions were selected for fragmentation with an isolation window of 4 and normalized HCD collision energy of 27%; the maximum injection time was set to 250 ms and AGC target was 5×10^5^ in an orbitrap mass analyzer with a resolution of 17,500 FWHM at m/z 200.

**MS instrument parameters for glycopeptide analysis**

MS parameters were set as follows: nano spray ion source was set at a capillary voltage of 1.8 kV. for MS1 scanning, m/z scan range was set from 350 to 2,000; an orbitrap mass analyzer was used with an orbitrap resolution of 120,000 FWHM at m/z 200; the maximum injection time was 50 ms and AGC target was 5×10^5^. Using top speed mode, HCD-MS/MS was performed with an isolation window of 4; isolated precursor ions were fragmented by a stepped HCD gradient with normalized collision energies of 20%, 30% and 40%; fragment ions were analyzed in an orbitrap mass analyzer at an orbitrap resolution of 15,000 FWHM at m/z 200; the maximum injection time was 250 ms and AGC target was 5×10^5^.

**MS instrument parameters for de-N-glycosylated EPO analysis**

The MS parameters were listed as follows: electrospray ionization source was set at a spray voltage of 3.5 kV; in-source CID was set to 20 eV; precursor ions were accumulated for a maximum injection time 240 ms and AGC target was 3×10^6^; the m/z range was set to 750 to 4,000; ions were analyzed at the MS1 level using an orbitrap mass analyzer at an orbitrap resolution of 140,000 FWHM at m/z 200.

**Table Captions**

Table S1. The identified 52 N-glycan compositions (72 N-glycan structures) from A2F N-glycan standards at MS1 and MS2 levels.

Table S2. The identified 140 N-glycan compositions (237 N-glycan structures) from EPO at MS1 and MS2 levels.

Table S3. The summary of tryptic glycopeptide analysis by pGlyco software.

Table S4. The summary of chymotryptic glycopeptide analysis by pGlyco software.

Table S5. The identified glycopeptides by chymotrypsin digestion and comparison with permethylated N-glycan analysis.

**Figure Captions**

Figure S1. The newly designed Python script for the identification of permethylated N-glycans, considering trimannosylchitobiose core.

Figure S2. The additionally designed Python script, especially for the identification of permethylated N-glycans containing phosphorylated Man.

Figure S3. The characterization of Fuc_1_Red-HexNAc_1_, by EIC and MS2 fragmentation from N-glycans of EPO.

Figure S4. The identified peptides of EPO-3 based on bottom-up analysis after trypsin and chymotrypsin digestion by pFind software, with the known modifications.

Figure S5. The characterization of GAQKEAISPPDAAS(126)AAPL at MS2 level from chymotryptic digestion.

Figure S6. The comparison of N-glycan compositions identified from EPO-3 at different levels, including glycopeptide analysis using pGlyco software and permethylated N-glycan analysis by the Python scripts.

Table S1. The identified 52 N-glycan compositions (72 N-glycan structures) from A2F N-glycan standards at MS1 and MS2 levels.

| **No.** | **N-glycan composition** | **No. of isomeric structures** | **Retention time (min)** | **Preferred N-glycan structure** | **Permethylation monoisotopic MW (Da)** | | **Deviation (p.p.m.)** |
| --- | --- | --- | --- | --- | --- | --- | --- |
|  |  |  |  |  | **Experiment** | **Theory** |  |
| 1 | Fuc_1_Red-HexNAc_1_ | 1 | 35.87 | 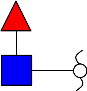 | 481.28865 | 481.2887 | 0.10 |
| 2 | Hex_1_Red-HexNAc_1_ | 1 | 37.47 | - | 511.29913 | 511.2993 | 0.25 |
| 3 | HexNAc_1_Fuc_1_Red-HexNAc_1_ | 1 | 40.79 | 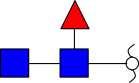 | 726.41519 | 726.4153 | 0.23 |
| 4 | Neu5Ac_1_Hex_1_Red-HexNAc_1_ | 1 | 43.31 | - | 872.47347 | 872.4729 | 0.63 |
| 5 | HexNAc_2_Hex_3_Red-HexNAc_1_ | 1 | 50.01 | 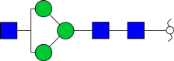 | 1409.7519 | 1409.7515 | 0.33 |
| 6 | Neu5Ac_1_HexNAc_1_Hex_3_Red-HexNAc_1_ | 2 | 52.76, 53.46 | 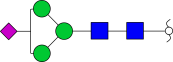 | 1525.7985 | 1525.7988 | 0.18 |
| 7 | HexNAc_1_Hex_5_Red-HexNAc_1_ | 1 | 56.83 | 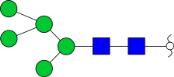 | 1572.8244 | 1572.8247 | 0.17 |
| 8 | HexNAc_2_Hex_3_Fuc_1_Red-HexNAc_1_ | 3 | 51.47, 52.25, 53.51 | 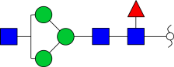 | 1583.8405 | 1583.8407 | 0.09 |
| 9 | HexNAc_2_Hex_4_Red-HexNAc_1_ | 2 | 52.38, 54.74 | 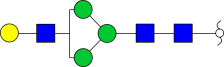 | 1613.8498 | 1613.8512 | 0.87 |
|  |  |  |  | 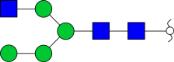 |  |  |  |
| 10 | HexNAc_3_Hex_3_Red-HexNAc_1_ | 1 | 52.19 | 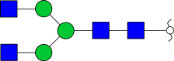 | 1654.8776 | 1654.8778 | 0.10 |
| 11 | Neu5Ac_1_HexNAc_2_Hex_3_Red-HexNAc_1_ | 1 | 53.10 | 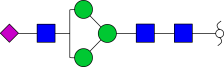 | 1770.9241 | 1770.9251 | 0.56 |
| 12 | HexNAc_1_Hex_6_Red-HexNAc_1_ | 1 | 61.16 | 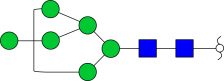 | 1776.9247 | 1776.9245 | 0.15 |
| 13 | HexNAc_2_Hex_4_Fuc_1_Red-HexNAc_1_ | 1 | 55.22 | 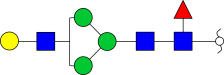 | 1787.9401 | 1787.9404 | 0.18 |
| 14 | HexNAc_2_Hex_5_Red-HexNAc_1_ | 2 | 56.47, 56.99 | 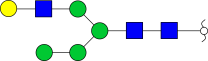 | 1817.9508 | 1817.9510 | 0.10 |
| 15 | HexNAc_3_Hex_3_Fuc_1_Red-HexNAc_1_ | 2 | 54.99, 55.60 | 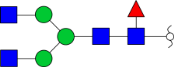 | 1828.9669 | 1828.9670 | 0.04 |
| 16 | HexNAc_3_Hex_4_Red-HexNAc_1_ | 1 | 53.83 | 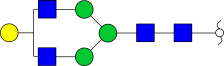 | 1858.9789 | 1858.9776 | 0.74 |
| 17 | Neu5Ac_1_HexNAc_2_Hex_3_Fuc_1_Red-HexNAc_1_ | 1 | 55.65 | 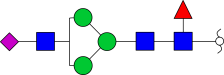 | 1945.0135 | 1945.0143 | 0.42 |
| 18 | HexNAc_3_Hex_4_Fuc_1_Red-HexNAc_1_ | 1 | 56.40 | 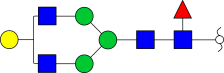 | 2033.065 | 2033.0668 | 0.86 |
| 19 | HexNAc_3_Hex_5_Red-HexNAc_1_ | 1 | 55.31 | 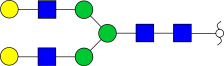 | 2063.077 | 2063.0773 | 0.15 |
| 20 | Neu5Ac_1_HexNAc_2_Hex_4_Fuc_1_Red-HexNAc_1_ | 1 | 58.24 | 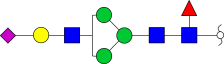 | 2149.1145 | 2149.1141 | 0.20 |
| 21 | Neu5Ac_1_HexNAc_2_Hex_5_Red-HexNAc_1_ | 1 | 59.13 | 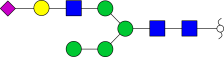 | 2179.1248 | 2179.1247 | 0.07 |
| 22 | HexNAc_1_Hex_8_Red-HexNAc_1_ | 2 | 67.87, 68.29 | 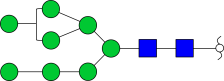 | 2185.1242 | 2185.1240 | 0.11 |
| 23 | Neu5Ac_1_HexNAc_3_Hex_4_Red-HexNAc_1_ | 1 | 56.84 | 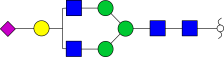 | 2220.1503 | 2220.1512 | 0.40 |
| 24 | HexNAc_3_Hex_5_Fuc_1_Red-HexNAc_1_ | 1 | 57.84 | 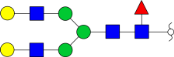 | 2237.1669 | 2237.1665 | 0.17 |
| 25 | Neu5Gc_1_HexNAc_3_Hex_4_Red-HexNAc_1_ | 1 | 56.29 | 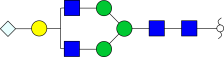 | 2250.1607 | 2250.1618 | 0.47 |
| 26 | Neu5Ac_1_HexNAc_3_Hex_3_Fuc_2_Red-HexNAc_1_ | 1 | 57.39 | 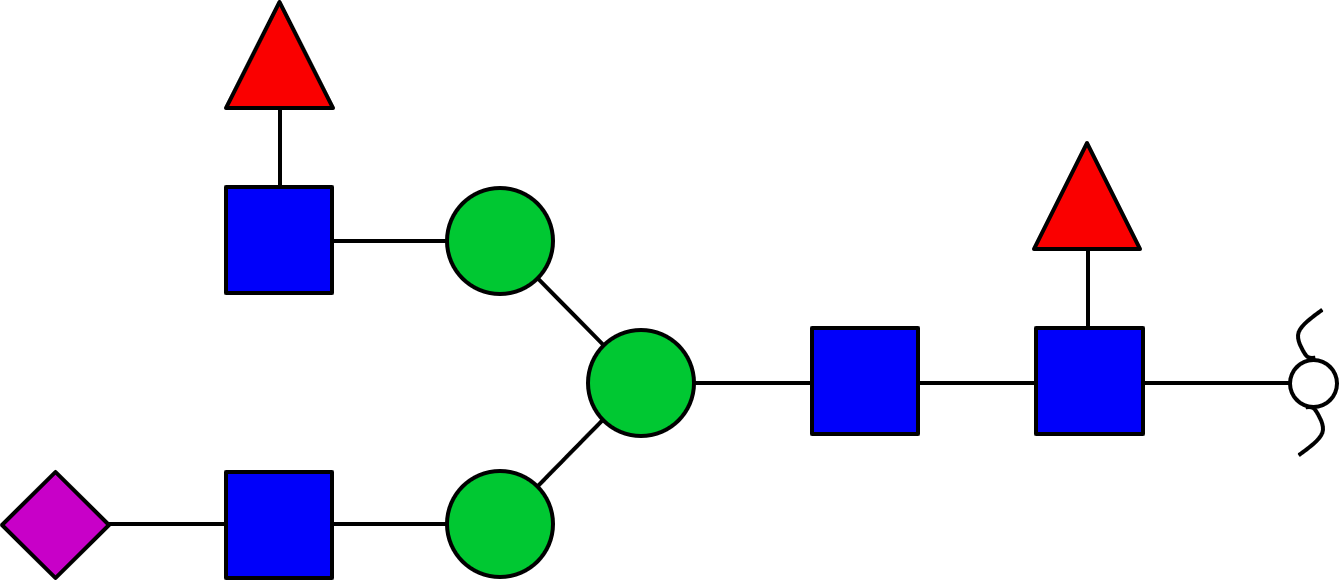 | 2364.2289 | 2364.2299 | 0.40 |
| 27 | HexNAc_1_Hex_9_Red-HexNAc_1_ | 1 | 70.95 | 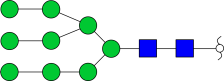 | 2389.229 | 2389.2238 | 2.20 |
| 28 | Neu5Ac_1_HexNAc_3_Hex_4_Fuc_1_Red-HexNAc_1_ | 1 | 58.98 | 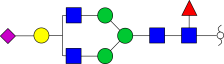 | 2394.2406 | 2394.2404 | 0.08 |
| 29 | Neu5Ac_1_HexNAc_3_Hex_5_Red-HexNAc_1_ | 3 | 58.01, 58.42, 59.23 | 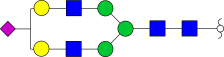 | 2424.2518 | 2424.2510 | 0.35 |
| 30 | Neu5Gc_1_HexNAc_3_Hex_4_Fuc_1_Red-HexNAc_1_ |  |  | 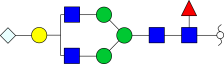 |  |  |  |
| 31 | Neu5Gc_1_HexNAc_3_Hex_5_Red-HexNAc_1_ | 1 | 57.44 | 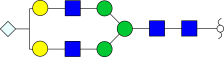 | 2454.2606 | 2454.2616 | 0.38 |
| 32 | Neu5Ac_2_HexNAc_2_Hex_5_Red-HexNAc_1_ | 2 | 61.15, 63.14 | 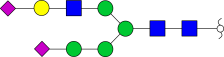 | 2540.2973 | 2540.2983 | 0.39 |
| 33 | HexNAc_1_Hex_10_Red-HexNAc_1_ | 1 | 76.67 | 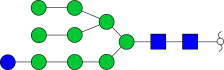 | 2593.3254 | 2593.3235 | 0.73 |
| 34 | Neu5Ac_1_HexNAc_3_Hex_5_Fuc_1_Red-HexNAc_1_ | 1 | 60.02 | 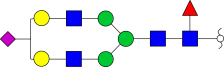 | 2598.3407 | 2598.3402 | 0.20 |
| 35 | Neu5Ac_1_HexNAc_3_Hex_6_Red-HexNAc_1_ | 1 | 59.56 | 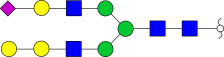 | 2628.3501 | 2628.3508 | 0.24 |
| 36 | HexNAc_5_Hex_6_Red-HexNAc_1_ | 2 | 37.85, 39.40 | 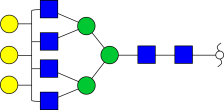 | 2757.4308 | 2757.4297 | 0.40 |
| 37 | Neu5Ac_2_HexNAc_3_Hex_5_Red-HexNAc_1_ | 1 | 59.82 | 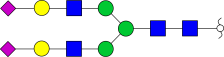 | 2785.4240 | 2785.4247 | 0.22 |
| 38 | Neu5Gc_1_Neu5Ac_1_HexNAc_3_Hex_5_Red-HexNAc_1_ | 1 | 59.31 | 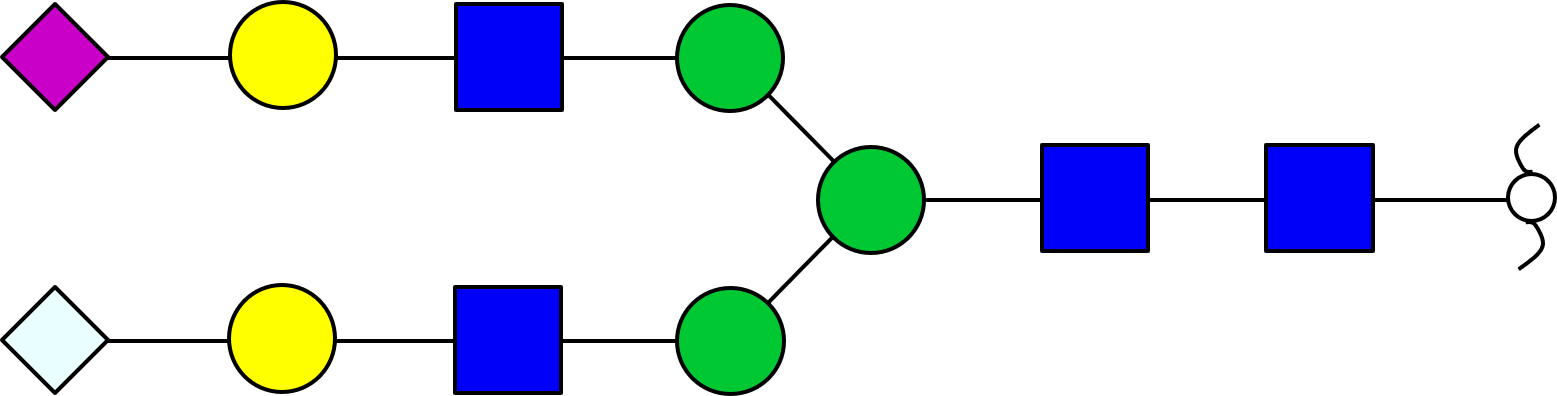 | 2815.4366 | 2815.4352 | 0.50 |
| 39 | Neu5Gc_2_HexNAc_3_Hex_5_Red-HexNAc_1_ | 2 | 58.86, 60.10 | 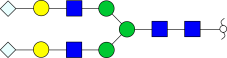 | 2845.4471 | 2845.4458 | 0.47 |
| 40 | HexNAc_5_Hex_6_Fuc_1_Red-HexNAc_1_ | 3 | 39.30, 41.37, 42.42 | 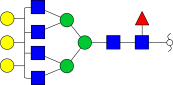 | 2931.5204 | 2931.5190 | 0.50 |
| 41 | Neu5Ac_2_HexNAc_3_Hex_5_Fuc_1_Red-HexNAc_1_ | 2 | 61.52, 65.78 | 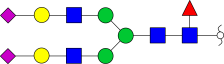 | 2959.5143 | 2959.5139 | 0.16 |
| 42 | Neu5Ac_1_HexNAc_4_Hex_4_Fuc_3_Red-HexNAc_1_ | 1 | 71.28 | 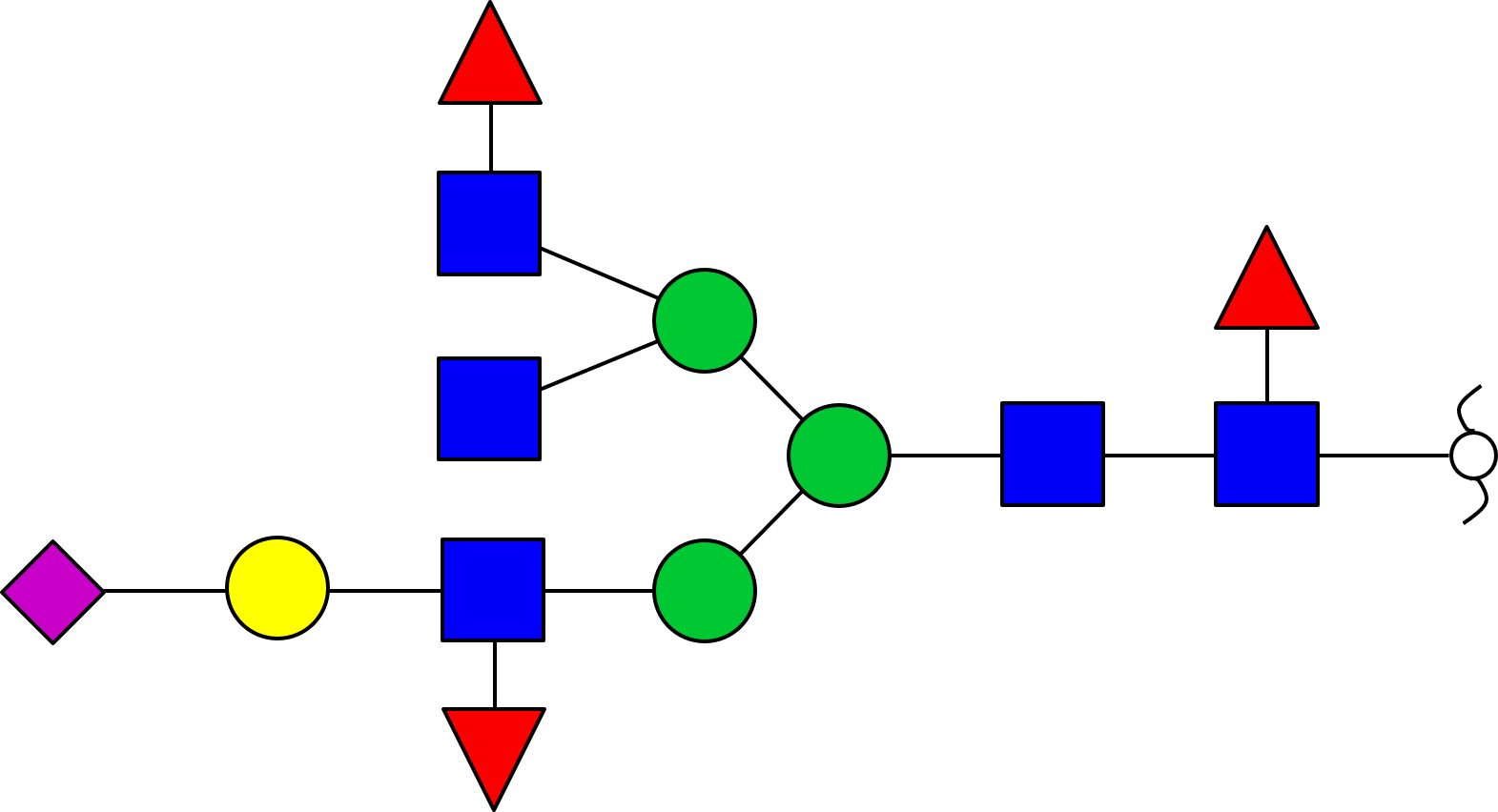 | 2987.5448 | 2987.5452 | 0.11 |
| 43 | Neu5Gc_1_Neu5Ac_1_HexNAc_3_Hex_5_Fuc_1_Red-HexNAc_1_ | 2 | 61.15, 62.15 | 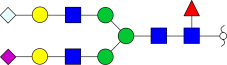 | 2989.5271 | 2989.5244 | 0.90 |
| 44 | Neu5Gc_1_HexNAc_3_Hex_6_Fuc_2_Red-HexNAc_1_ | 1 | 61.11 | 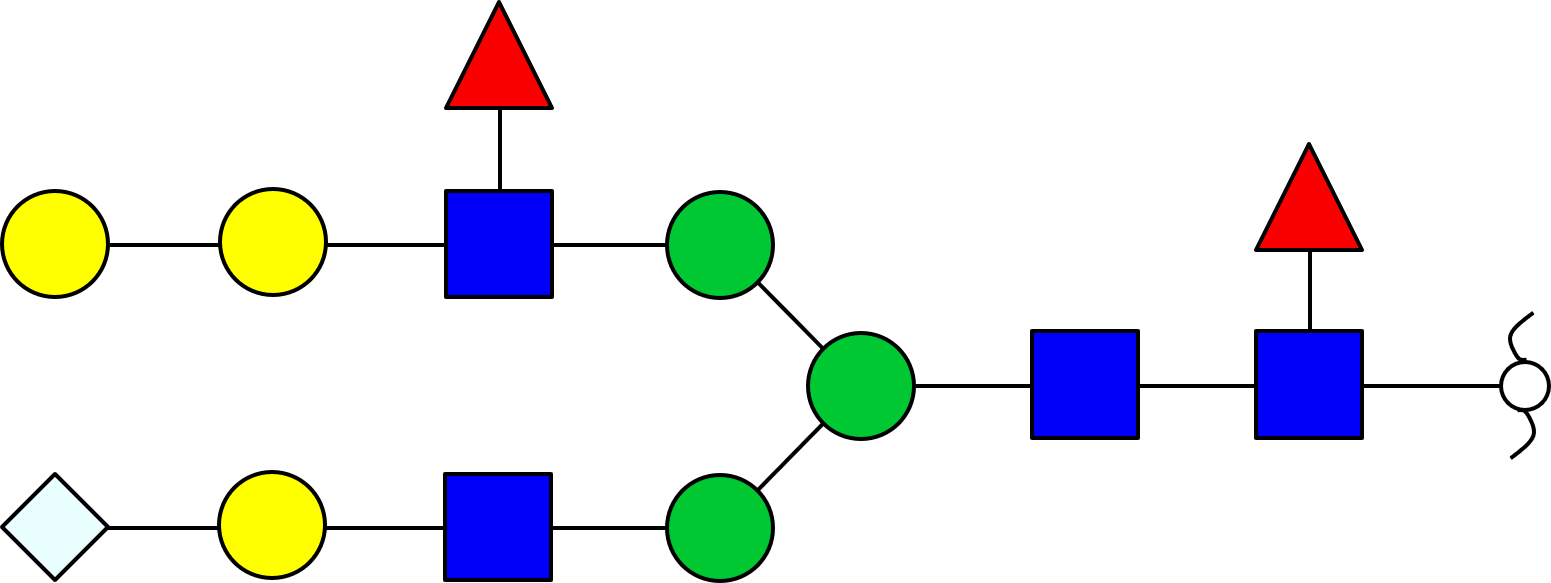 | 3006.5397 | 3006.5397 | 0.01 |
| 45 | Neu5Gc_1_Neu5Ac_1_HexNAc_3_Hex_6_Red-HexNAc_1_ | 2 | 60.70, 62.05 | 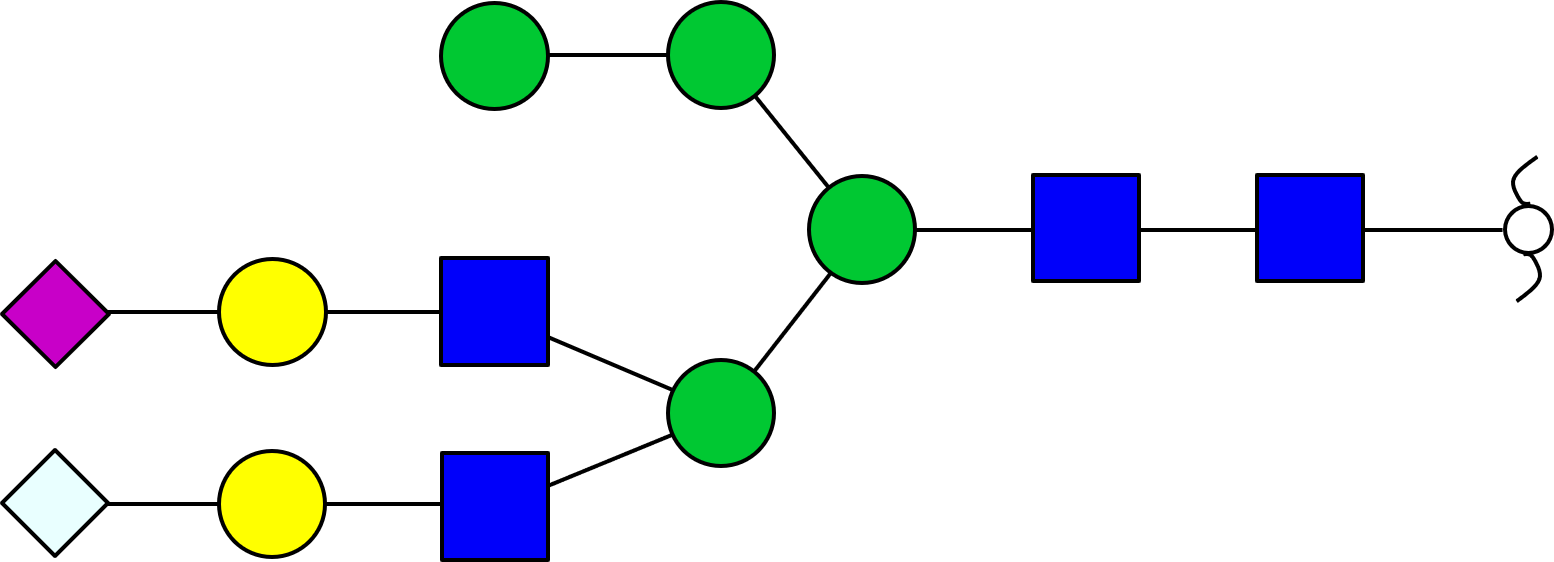 | 3019.534 | 3019.5350 | 0.32 |
| 46 | Neu5Ac_1_HexNAc_4_Hex_6_Fuc_1_Red-HexNAc_1_ | 1 | 59.92 | 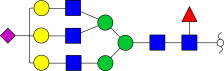 | 3047.565 | 3047.5663 | 0.41 |
| 47 | Neu5Ac_2_HexNAc_3_Hex_6_Fuc_1_Red-HexNAc_1_ | 3 | 63.90, 64.81, 66.24 | 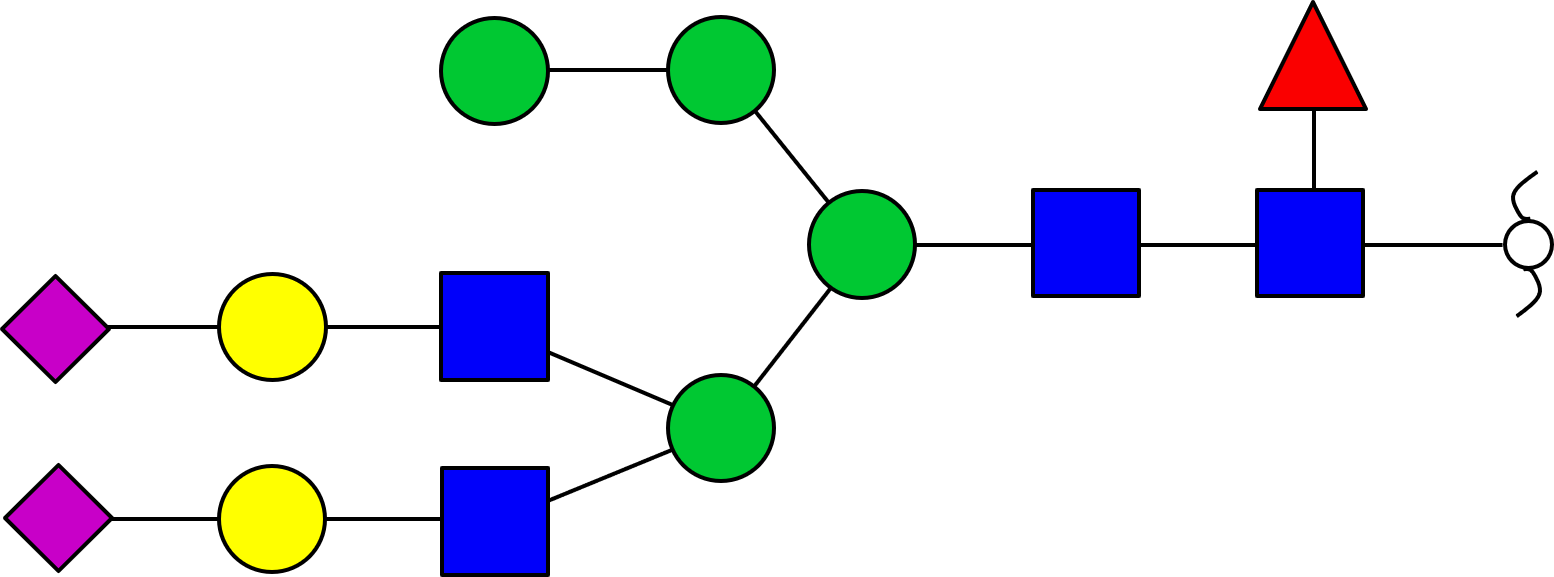 | 3163.6143 | 3163.6136 | 0.22 |
| 48 | Neu5Ac_2_HexNAc_4_Hex_5_Fuc_1_Red-HexNAc_1_ | 1 | 60.76 | 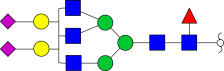 | 3204.6411 | 3204.6402 | 0.30 |
| 49 | Neu5Ac_2_HexNAc_4_Hex_6_Red-HexNAc_1_ | 1 | 60.35 | 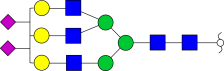 | 3234.6489 | 3234.6508 | 0.56 |
| 50 | Neu5Ac_3_HexNAc_3_Hex_5_Fuc_1_Red-HexNAc_1_ | 1 | 65.18 | 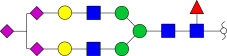 | 3320.689 | 3320.6875 | 0.45 |
| 51 | Neu5Ac_2_HexNAc_3_Hex_6_Fuc_2_Red-HexNAc_1_ | 1 | 65.17 | 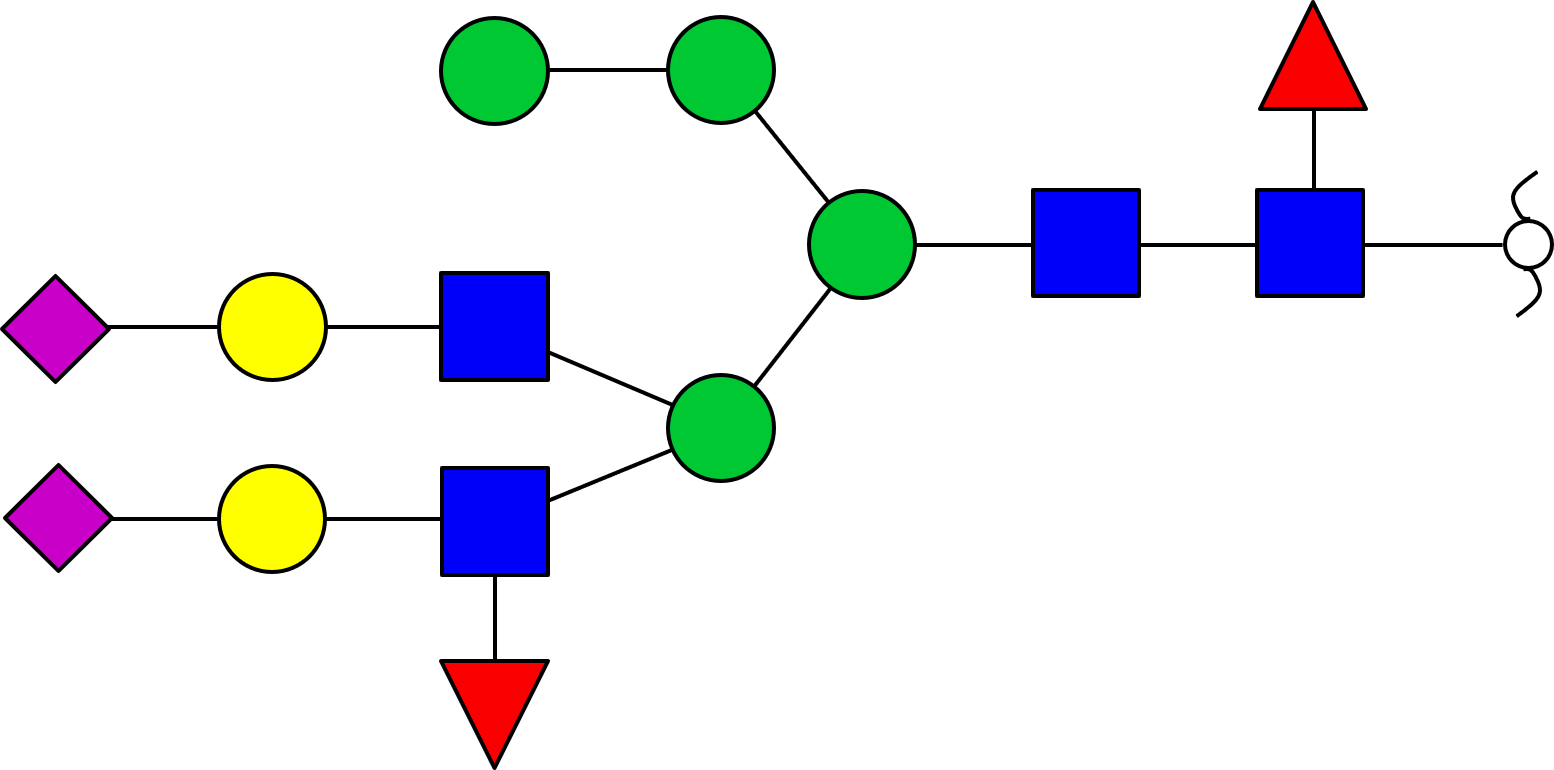 | 3337.7088 | 3337.7028 | 1.79 |
| 52 | Neu5Ac_2_HexNAc_4_Hex_6_Fuc_1_Red-HexNAc_1_ | 3 | 61.23, 62.10, 64.32 | 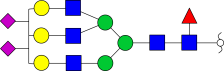 | 3408.7391 | 3408.7400 | 0.24 |

Table S2. The identified 140 N-glycan compositions (237 N-glycan structures) from EPO at MS1 and MS2 levels.

| **No.** | **N-glycan composition** | **No. of isomeric structures** | **Retention time (min)** | **Preferred N-glycan structure** | **Permethylation monoisotopic MW (Da)** | | **Deviation (p.p.m.)** |
| --- | --- | --- | --- | --- | --- | --- | --- |
|  |  |  |  |  | **Experiment** | **Theory** |  |
| 1 | Fuc_1_Red-HexNAc_1_ | 1 | 35.82 | 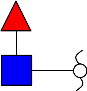 | 481.2888 | 481.2887 | 0.21 |
| 2 | Hex_1_Red-HexNAc_1_ | 3 | 37.42, 38.11, 38.84 | - | 511.29932 | 511.2993 | 0.12 |
| 3 | HexNAc_1_Fuc_1_Red-HexNAc_1_ | 1 | 40.77 | 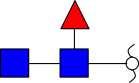 | 726.41529 | 726.4153 | 0.37 |
| 4 | Neu5Ac_1_Hex_1_Red-HexNAc_1_ | 3 | 46.19, 47.76, 49.51 | - | 872.47304 | 872.4729 | 0.14 |
| 5 | Neu5Gc_1_Hex_1_Red-HexNAc_1_ | 2 | 45.41, 47.76 | - | 902.48364 | 902.4835 | 0.17 |
| 6 | HexNAc_1_Hex_4_Red-HexNAc_1_ | 1 | 53.00 | 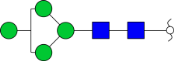 | 1368.7247 | 1368.7249 | 0.14 |
| 7 | HexNAc_2_Hex_3_Red-HexNAc_1_ | 1 | 48.02 | 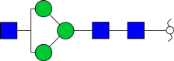 | 1409.7525 | 1409.7515 | 0.75 |
| 8 | Neu5Ac_1_HexNAc_1_Hex_3_Red-HexNAc_1_ | 1 | 54.65 | 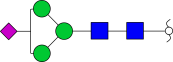 | 1525.7987 | 1525.7988 | 0.05 |
| 9 | HexNAc_1_Hex_4_Fuc_1_Red-HexNAc_1_ | 1 | 56.36 | 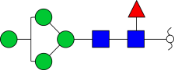 | 1542.8141 | 1542.8141 | 0.00 |
| 10 | HexNAc_1_Hex_5_Red-HexNAc_1_ | 1 | 56.85 | 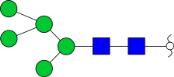 | 1572.8244 | 1572.8247 | 0.17 |
| 11 | HexNAc_2_Hex_3_Fuc_1_Red-HexNAc_1_ | 3 | 51.53, 52.38, 53.52 | 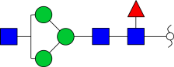 | 1583.8407 | 1583.8407 | 0.03 |
| 12 | HexNAc_2_Hex_4_Red-HexNAc_1_ | 1 | 52.38 | 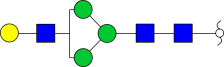 | 1613.8519 | 1613.8512 | 0.42 |
| 13 | HexNAc_3_Hex_3_Red-HexNAc_1_ | 1 | 52.18 | 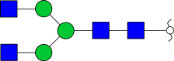 | 1654.8796 | 1654.8778 | 1.11 |
| 14 | HexNAc_1_Hex_6_Red-HexNAc_1_ | 1 | 60.85 | 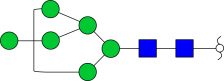 | 1776.9253 | 1776.9245 | 0.49 |
| 15 | HexNAc_2_Hex_4_Fuc_1_Red-HexNAc_1_ | 1 | 55.19 | 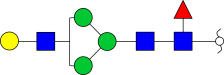 | 1787.9406 | 1787.9404 | 0.10 |
| 16 | HexNAc_3_Hex_3_Fuc_1_Red-HexNAc_1_ | 1 | 54.96 | 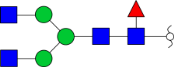 | 1828.9672 | 1828.9670 | 0.13 |
| 17 | HexNAc_3_Hex_4_Red-HexNAc_1_ | 1 | 53.82 | 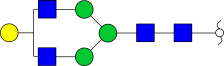 | 1858.9786 | 1858.9776 | 0.58 |
| 18 | Neu5Ac_1_HexNAc_2_Hex_3_Fuc_1_Red-HexNAc_1_ | 1 | 56.58 | 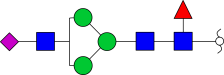 | 1945.0159 | 1945.0143 | 0.82 |
| 19 | Neu5Gc_1_HexNAc_2_Hex_4_Red-HexNAc_1_ | 1 | 52.18 | 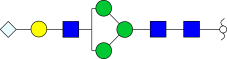 | 2005.0362 | 2005.0355 | 0.38 |
| 20 | HexNAc_3_Hex_4_Fuc_1_Red-HexNAc_1_ | 1 | 56.48 | 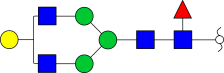 | 2033.0667 | 2033.0668 | 0.02 |
| 21 | Neu5Ac_1_HexNAc_2_Hex_4_Fuc_1_Red-HexNAc_1_ | 1 | 58.84 | 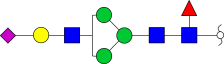 | 2149.1130 | 2149.1141 | 0.50 |
| 22 | HexNAc_3_Hex_5_Fuc_1_Red-HexNAc_1_ | 1 | 57.83 | 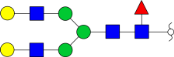 | 2237.1677 | 2237.1665 | 0.53 |
| 23 | Neu5Ac_1_HexNAc_3_Hex_4_Fuc_1_Red-HexNAc_1_ | 2 | 58.84, 59.58 | 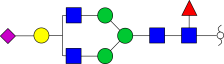 | 2394.2397 | 2394.2404 | 0.29 |
| 24 | Neu5Gc_1_HexNAc_3_Hex_5_Red-HexNAc_1_ | 1 | 57.35 | 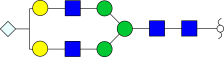 | 2454.2604 | 2454.2616 | 0.46 |
| 25 | Neu5Ac_1_HexNAc_3_Hex_4_Fuc_2_Red-HexNAc_1_ | 2 | 57.47, 59.64 | 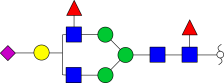 | 2568.3296 | 2568.3296 | 0.00 |
| 26 | Neu5Ac_1_HexNAc_3_Hex_5_Fuc_1_Red-HexNAc_1_ | 2 | 58.45, 60.95 | 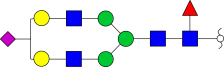 | 2598.3408 | 2598.3402 | 0.24 |
| 27 | Neu5Gc_1_HexNAc_3_Hex_5_Fuc_1_Red-HexNAc_1_ | 2 | 59.37, 60.23 | 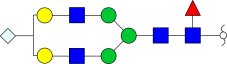 | 2628.3504 | 2628.3508 | 0.13 |
| 28 | Neu5Ac_2_HexNAc_3_Hex_3_Fuc_2_Red-HexNAc_1_ | 1 | 58.72 | 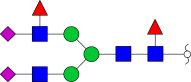 | 2725.4019 | 2725.4035 | 0.59 |
| 29 | Neu5Ac_2_HexNAc_3_Hex_4_Fuc_1_Red-HexNAc_1_ | 2 | 60.22, 61.68 | 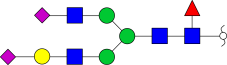 | 2755.4143 | 2755.4141 | 0.09 |
| 30 | Neu5Ac_1_HexNAc_2_Hex_7_Fuc_1_Red-HexNAc_1_ | 1 | 69.85 | 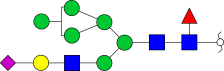 | 2761.4146 | 2761.4134 | 0.44 |
| 31 | Neu5Ac_1_HexNAc_4_Hex_5_Fuc_1_Red-HexNAc_1_ | 1 | 59.81 | 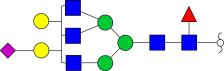 | 2843.4652 | 2843.4665 | 0.45 |
| 32 | Neu5Ac_2_HexNAc_3_Hex_4_Fuc_2_Red-HexNAc_1_ | 3 | 60.03, 61.38, 62.00 | 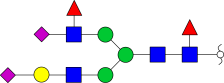 | 2929.5057 | 2929.5033 | 0.83 |
| 33 | Neu5Ac_2_HexNAc_3_Hex_5_Fuc_1_Red-HexNAc_1_ | 1 | 61.61 | 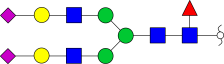 | 2959.5142 | 2959.5139 | 0.13 |
| 34 | Neu5Gc_1_Neu5Ac_1_HexNAc_3_Hex_5_Fuc_1_Red-HexNAc_1_ | 3 | 61.25, 64.25, 63.30 | 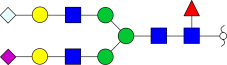 | 2989.5234 | 2989.5244 | 0.33 |
| 35 | Neu5Ac_1_HexNAc_4_Hex_6_Fuc_1_Red-HexNAc_1_ | 2 | 60.57, 62.58 | 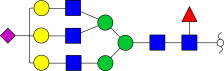 | 3047.5661 | 3047.5663 | 0.05 |
| 36 | Neu5Gc_1_HexNAc_4_Hex_6_Fuc_1_Red-HexNAc_1_ | 2 | 60.23, 61.96 | 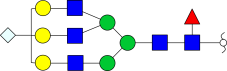 | 3077.5776 | 3077.5769 | 0.25 |
| 37 | Neu5Ac_3_HexNAc_3_Hex_5_Fuc_1_Red-HexNAc_1_ | 1 | 64.47 | 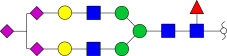 | 3320.6882 | 3320.6875 | 0.21 |
| 38 | Neu5Ac_2_HexNAc_4_Hex_5_Fuc_2_Red-HexNAc_1_ | 1 | 61.31 | 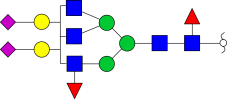 | 3378.7254 | 3378.7294 | 1.17 |
| 39 | Neu5Gc_1_Neu5Ac_2_HexNAc_3_Hex_6_Red-HexNAc_1_ | 3 | 56.64, 60.06, 62.40 | 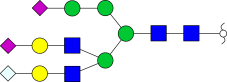 | 3380.7086 | 3380.7087 | 0.01 |
| 40 | HexNAc_6_Hex_7_Fuc_1_Red-HexNAc_1_ | 1 | 42.94 | 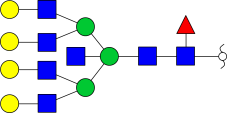 | 3380.7434 | 3380.7450 | 0.48 |
| 41 | Neu5Ac_2_HexNAc_4_Hex_6_Fuc_1_Red-HexNAc_1_ | 2 | 62.83, 64.21 | 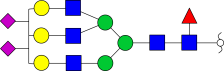 | 3408.7409 | 3408.7400 | 0.29 |
| 42 | Neu5Gc_1_Neu5Ac_1_HexNAc_4_Hex_6_Fuc_1_Red-HexNAc_1_ | 2 | 62.40, 63.87 | 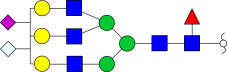 | 3438.7502 | 3438.7505 | 0.08 |
| 43 | Neu5Gc_1_Neu5Ac_1_HexNAc_4_Hex_7_Red-HexNAc_1_ | 2 | 53.32, 56.70 | 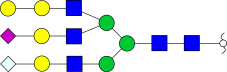 | 3468.7621 | 3468.7611 | 0.30 |
| 44 | Neu5Ac_3_HexNAc_4_Hex_6_Red-HexNAc_1_ | 1 | 63.06 | 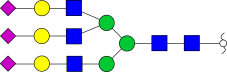 | 3595.8228 | 3595.8244 | 0.44 |
| 45 | Neu5Gc_1_Neu5Ac_2_HexNAc_4_Hex_6_Red-HexNAc_1_ | 2 | 58.82, 63.51 | 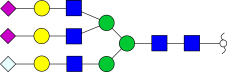 | 3625.8343 | 3625.8350 | 0.18 |
| 46 | Neu5Ac_2_HexNAc_5_Hex_6_Fuc_1_Red-HexNAc_1_ | 1 | 63.44 | 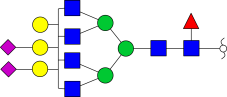 | 3653.8652 | 3653.8663 | 0.28 |
| 47 | Neu5Gc_1_Neu5Ac_3_HexNAc_3_Hex_5_Fuc_1_Red-HexNAc_1_ | 1 | 61.09 | 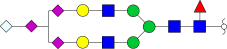 | 3711.8718 | 3711.8718 | 0.02 |
| 48 | Neu5Gc_1_Neu5Ac_1_HexNAc_5_Hex_7_Red-HexNAc_1_ | 5 | 41.47, 42.02, 42.50, 43.30, 43.84 | 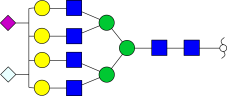 | 3713.8878 | 3713.8874 | 0.12 |
| 49 | Neu5Ac_3_HexNAc_4_Hex_5_Fuc_2_Red-HexNAc_1_ | 3 | 62.02, 63.25, 65.61 | 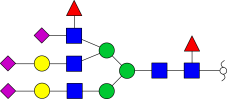 | 3739.9054 | 3739.9031 | 0.64 |
| 50 | Neu5Gc_2_Neu5Ac_2_HexNAc_3_Hex_5_Fuc_1_Red-HexNAc_1_ | 2 | 60.44, 61.10 | 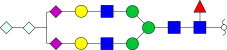 | 3741.8843 | 3741.8823 | 0.54 |
| 51 | Neu5Ac_1_HexNAc_6_Hex_7_Fuc_1_Red-HexNAc_1_ | 2 | 45.19, 46.21 | 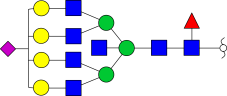 | 3741.9187 | 3741.9187 | 0.01 |
| 52 | Neu5Ac_3_HexNAc_4_Hex_6_Fuc_1_Red-HexNAc_1_ | 2 | 64.54, 65.89 | 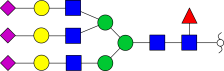 | 3769.9144 | 3769.9136 | 0.22 |
| 53 | Neu5Ac_1_HexNAc_7_Hex_6_Fuc_1_Red-HexNAc_1_ | 6 | 39.96, 40.80, 41.31, 41.89, 42.68, 44.68 |  | 3782.9441 | 3782.9453 | 0.30 |
| 54 | Neu5Ac_2_HexNAc_4_Hex_7_Fuc_2_Red-HexNAc_1_ | 2 | 64.60, 65.82 |  | 3786.9284 | 3786.9289 | 0.13 |
| 55 | Neu5Gc_1_Neu5Ac_2_HexNAc_4_Hex_6_Fuc_1_Red-HexNAc_1_ | 2 | 64.26, 65.60 |  | 3799.9242 | 3799.9242 | 0.02 |
| 56 | Neu5Ac_1_HexNAc_7_Hex_7_Red-HexNAc_1_ | 2 | 39.94, 41.96 |  | 3812.9570 | 3812.9558 | 0.32 |
| 57 | Neu5Gc_1_Neu5Ac_1_HexNAc_4_Hex_7_Fuc_2_Red-HexNAc_1_ | 2 | 64.28, 65.50 |  | 3816.9393 | 3816.9395 | 0.04 |
| 58 | Neu5Gc_2_Neu5Ac_1_HexNAc_4_Hex_6_Fuc_1_Red-HexNAc_1_ | 2 | 59.41, 62.18 |  | 3829.9346 | 3829.9348 | 0.03 |
| 59 | Neu5Ac_2_HexNAc_5_Hex_7_Fuc_1_Red-HexNAc_1_ | 2 | 64.38, 65.64 |  | 3857.9663 | 3857.9661 | 0.08 |
| 60 | Neu5Ac_1_HexNAc_5_Hex_8_Fuc_2_Red-HexNAc_1_ | 2 | 64.35, 65.79 |  | 3874.9905 | 3874.9814 | 2.37 |
| 61 | Neu5Gc_1_Neu5Ac_1_HexNAc_5_Hex_7_Fuc_1_Red-HexNAc_1_ | 2 | 60.63, 66.37 |  | 3887.9765 | 3887.9766 | 0.02 |
| 62 | Neu5Gc_1_Neu5Ac_1_HexNAc_5_Hex_8_Red-HexNAc_1_ | 1 | 66.50 |  | 3917.9884 | 3917.9872 | 0.32 |
| 63 | Neu5Ac_3_HexNAc_4_Hex_6_Fuc_2_Red-HexNAc_1_ | 1 | 66.81 |  | 3944.0018 | 3944.0028 | 0.25 |
| 64 | Neu5Ac_3_HexNAc_5_Hex_6_Fuc_1_Red-HexNAc_1_ | 1 | 65.30 |  | 4015.0397 | 4015.0400 | 0.05 |
| 65 | Neu5Ac_2_HexNAc_5_Hex_7_Fuc_2_Red-HexNAc_1_ | 1 | 65.59 |  | 4032.0626 | 4032.0553 | 1.83 |
| 66 | Neu5Ac_2_HexNAc_6_Hex_7_Fuc_1_Red-HexNAc_1_ | 1 | 64.44 |  | 4103.0926 | 4103.0924 | 0.07 |
| 67 | Neu5Ac_4_HexNAc_4_Hex_6_Fuc_1_Red-HexNAc_1_ | 2 | 65.49, 67.10 |  | 4131.0870 | 4131.0873 | 0.06 |
| 68/  69 | Neu5Gc_1_Neu5Ac_3_HexNAc_4_Hex_6_Fuc_1_Red-HexNAc_1_ | 3 | 59.21, 63.69 |  | 4161.0981 | 4161.0979 | 0.07 |
|  | Neu5Ac_4_HexNAc_4_Hex_7_Red-HexNAc_1_ |  | 67.52 |  |  |  |  |
| 70 | Neu5Gc_4_Neu5Ac_1_HexNAc_3_Hex_5_Fuc_1_Red-HexNAc_1_ | 2 | 55.52, 58.74 |  | 4163.0780 | 4163.0771 | 0.22 |
| 71 | Neu5Gc_1_Neu5Ac_1_HexNAc_6_Hex_8_Red-HexNAc_1_ | 5 | 42.80, 43.40, 43.92, 44.62, 45.74 |  | 4163.1148 | 4163.1135 | 0.32 |
| 72 | Neu5Ac_3_HexNAc_5_Hex_6_Fuc_2_Red-HexNAc_1_ | 2 | 63.53, 64.90 |  | 4189.1253 | 4189.1292 | 0.91 |
| 73 | Neu5Gc_2_Neu5Ac_2_HexNAc_4_Hex_6_Fuc_1_Red-HexNAc_1_ | 2 | 61.84, 64.35 |  | 4191.1097 | 4191.1084 | 0.32 |
| 74 | Neu5Ac_1_HexNAc_7_Hex_8_Fuc_1_Red-HexNAc_1_ | 1 | 46.71 |  | 4191.1439 | 4191.1448 | 0.21 |
| 75 | Neu5Gc_1_Neu5Ac_1_HexNAc_7_Hex_7_Red-HexNAc_1_ | 1 | 37.52 |  | 4204.1420 | 4204.1401 | 0.47 |
| 76 | Neu5Ac_3_HexNAc_5_Hex_7_Fuc_1_Red-HexNAc_1_ | 3 | 65.55, 65.90, 66.80 |  | 4219.1407 | 4219.1397 | 0.24 |
| 77 | Neu5Gc_3_Neu5Ac_1_HexNAc_4_Hex_6_Fuc_1_Red-HexNAc_1_ | 1 | 61.27 |  | 4221.1171 | 4221.1190 | 0.44 |
| 78 | Neu5Ac_6_HexNAc_3_Hex_5_Red-HexNAc_1_ | 1 | 53.73 |  | 4230.1226 | 4230.1193 | 0.79 |
| 79 | Neu5Ac_1_HexNAc_8_Hex_7_Fuc_1_Red-HexNAc_1_ | 3 | 40.35, 41.70, 44.63 |  | 4232.1703 | 4232.1714 | 0.24 |
| 80 | Neu5Ac_2_HexNAc_5_Hex_8_Fuc_2_Red-HexNAc_1_ | 3 | 65.42, 65.90, 67.00 |  | 4236.1522 | 4236.1550 | 0.66 |
| 81 | Neu5Gc_1_Neu5Ac_2_HexNAc_5_Hex_7_Fuc_1_Red-HexNAc_1_ | 3 | 65.11, 65.61, 66.76 |  | 4249.1495 | 4249.1503 | 0.17 |
| 82 | Neu5Gc_1_Neu5Ac_1_HexNAc_5_Hex_8_Fuc_2_Red-HexNAc_1_ | 1 | 65.60 |  | 4266.1663 | 4266.1656 | 0.18 |
| 83 | Neu5Gc_2_Neu5Ac_1_HexNAc_5_Hex_7_Fuc_1_Red-HexNAc_1_ | 3 | 60.46, 63.83, 65.46 |  | 4279.1636 | 4279.1609 | 0.65 |
| 84 | Neu5Ac_2_HexNAc_6_Hex_8_Fuc_1_Red-HexNAc_1_ | 3 | 65.20, 65.68, 66.39 |  | 4307.1904 | 4307.1922 | 0.39 |
| 85 | Neu5Ac_1_HexNAc_6_Hex_9_Fuc_2_Red-HexNAc_1_ | 1 | 65.57 |  | 4324.2165 | 4324.2075 | 2.10 |
| 86 | Neu5Gc_1_Neu5Ac_3_HexNAc_5_Hex_7_Red-HexNAc_1_ | 1 | 65.51 |  | 4436.2375 | 4436.2347 | 0.63 |
| 87 | Neu5Ac_3_HexNAc_6_Hex_7_Fuc_1_Red-HexNAc_1_ | 1 | 66.09 |  | 4464.2622 | 4464.2660 | 0.85 |
| 88 | Neu5Ac_5_HexNAc_4_Hex_6_Fuc_1_Red-HexNAc_1_ | 1 | 66.98 |  | 4492.2600 | 4492.2610 | 0.20 |
| 89 | Neu5Ac_3_HexNAc_6_Hex_8_Red-HexNAc_1_ | 1 | 43.88 |  | 4494.2732 | 4494.2766 | 0.75 |
| 90 | Neu5Ac_4_HexNAc_5_Hex_6_Fuc_2_Red-HexNAc_1_ | 5 | 64.69, 65.75, 66.22, 66.64, 69.06 |  | 4550.2993 | 4550.3028 | 0.76 |
| 91 | Neu5Gc_2_Neu5Ac_3_HexNAc_4_Hex_6_Fuc_1_Red-HexNAc_1_ | 5 | 62.20, 62.77, 63.47, 63.63, 67.12 |  | 4552.2834 | 4552.2821 | 0.30 |
| 92 | Neu5Ac_2_HexNAc_7_Hex_8_Fuc_1_Red-HexNAc_1_ | 1 | 48.33 |  | 4552.3202 | 4552.3185 | 0.39 |
| 93/  94 | Neu5Gc_1_Neu5Ac_3_HexNAc_5_Hex_6_Fuc_2_Red-HexNAc_1_ | 4 | 64.53, 65.39 |  | 4580.3159 | 4580.3134 | 0.56 |
|  | Neu5Ac_4_HexNAc_5_Hex_7_Fuc_1_Red-HexNAc_1_ |  | 67.20, 68.20 |  |  |  |  |
| 95 | Neu5Gc_3_Neu5Ac_2_HexNAc_4_Hex_6_Fuc_1_Red-HexNAc_1_ | 1 | 63.00 |  | 4582.2919 | 4582.2927 | 0.15 |
| 96 | Neu5Ac_3_HexNAc_5_Hex_8_Fuc_2_Red-HexNAc_1_ | 1 | 67.09 |  | 4597.3299 | 4597.3287 | 0.27 |
| 97 | Neu5Gc_1_Neu5Ac_3_HexNAc_5_Hex_7_Fuc_1_Red-HexNAc_1_ | 3 | 67.00, 67.52, 68.50 |  | 4610.3204 | 4610.3240 | 0.76 |
| 98 | Neu5Ac_2_HexNAc_8_Hex_8_Red-HexNAc_1_ | 1 | 44.00 |  | 4623.3534 | 4623.3556 | 0.46 |
| 99 | Neu5Gc_1_Neu5Ac_2_HexNAc_5_Hex_8_Fuc_2_Red-HexNAc_1_ | 1 | 66.82 |  | 4627.3411 | 4627.3393 | 0.41 |
| 100 | Neu5Ac_3_HexNAc_6_Hex_7_Fuc_2_Red-HexNAc_1_ | 2 | 64.77, 66.01 |  | 4638.3560 | 4638.3553 | 0.17 |
| 101 | Neu5Gc_2_Neu5Ac_2_HexNAc_5_Hex_7_Fuc_1_Red-HexNAc_1_ | 3 | 62.18, 66.66, 67.50 |  | 4640.3381 | 4640.3345 | 0.78 |
| 102 | Neu5Ac_3_HexNAc_6_Hex_8_Fuc_1_Red-HexNAc_1_ | 4 | 65.73, 66.39, 66.85, 67.53 |  | 4668.3611 | 4668.3658 | 1.00 |
| 103 | Neu5Ac_1_HexNAc_9_Hex_8_Fuc_1_Red-HexNAc_1_ | 3 | 43.77, 45.86, 46.42 |  | 4681.3945 | 4681.3975 | 0.62 |
| 104 | Neu5Ac_2_HexNAc_6_Hex_9_Fuc_2_Red-HexNAc_1_ | 1 | 66.47 |  | 4685.3842 | 4685.3811 | 0.67 |
| 105 | Neu5Gc_1_Neu5Ac_2_HexNAc_6_Hex_8_Fuc_1_Red-HexNAc_1_ | 2 | 66.11, 66.61 |  | 4698.3762 | 4698.3764 | 0.03 |
| 106 | Neu5Ac_1_HexNAc_9_Hex_9_Red-HexNAc_1_ | 1 | 72.24 |  | 4711.4122 | 4711.4080 | 0.90 |
| 107 | Neu5Ac_4_HexNAc_5_Hex_7_Fuc_2_Red-HexNAc_1_ | 1 | 68.61 |  | 4754.4031 | 4754.4026 | 0.12 |
| 108 | Neu5Ac_2_HexNAc_7_Hex_9_Fuc_1_Red-HexNAc_1_ | 1 | 65.90 |  | 4756.4168 | 4756.4183 | 0.29 |
| 109 | Neu5Ac_1_HexNAc_7_Hex_10_Fuc_2_Red-HexNAc_1_ | 1 | 66.20 |  | 4773.4350 | 4773.4336 | 0.31 |
| 110 | Neu5Gc_1_Neu5Ac_3_HexNAc_5_Hex_7_Fuc_2_Red-HexNAc_1_ | 2 | 68.45, 69.51 |  | 4784.4133 | 4784.4132 | 0.04 |
| 111 | Neu5Ac_1_HexNAc_11_Hex_7_Red-HexNAc_1_ | 1 | 46.34 |  | 4793.4588 | 4793.4611 | 0.47 |
| 112 | Neu5Ac_4_HexNAc_5_Hex_7_Fuc_3_Red-HexNAc_1_ | 1 | 70.45 |  | 4928.4890 | 4928.4918 | 0.56 |
| 113 | Neu5Ac_4_HexNAc_6_Hex_7_Fuc_2_Red-HexNAc_1_ | 3 | 65.76, 66.78, 69.80 |  | 4999.5300 | 4999.5289 | 0.23 |
| 114 | Neu5Gc_2_Neu5Ac_3_HexNAc_5_Hex_7_Fuc_1_Red-HexNAc_1_ | 2 | 63.89, 68.55 |  | 5001.5141 | 5001.5082 | 1.20 |
| 115 | Neu5Ac_2_HexNAc_8_Hex_9_Fuc_1_Red-HexNAc_1_ | 1 | 49.77 |  | 5001.5453 | 5001.5446 | 0.16 |
| 116 | Neu5Ac_4_HexNAc_6_Hex_8_Fuc_1_Red-HexNAc_1_ | 2 | 67.30, 67.81 |  | 5029.5380 | 5029.5395 | 0.28 |
| 117 | Neu5Ac_2_HexNAc_9_Hex_8_Fuc_1_Red-HexNAc_1_ | 1 | 45.60 |  | 5042.5718 | 5042.5711 | 0.15 |
| 118 | Neu5Ac_3_HexNAc_6_Hex_9_Fuc_2_Red-HexNAc_1_ | 2 | 67.35, 67.84 |  | 5046.5565 | 5046.5548 | 0.35 |
| 119 | Neu5Gc_1_Neu5Ac_3_HexNAc_6_Hex_8_Fuc_1_Red-HexNAc_1_ | 2 | 67.03, 67.59 |  | 5059.5473 | 5059.5501 | 0.53 |
| 120 | Neu5Ac_3_HexNAc_7_Hex_9_Fuc_1_Red-HexNAc_1_ | 2 | 66.64, 67.17 |  | 5117.5925 | 5117.5919 | 0.13 |
| 121 | Neu5Ac_1_HexNAc_10_Hex_9_Fuc_1_Red-HexNAc_1_ | 1 | 47.74 |  | 5130.6236 | 5130.6236 | 0.02 |
| 122 | Neu5Gc_1_Neu5Ac_1_HexNAc_7_Hex_9_Fuc_3_Red-HexNAc_1_ | 1 | 66.80 |  | 5134.6077 | 5134.6072 | 0.10 |
| 123 | Neu5Ac_1_HexNAc_10_Hex_10_Red-HexNAc_1_ | 1 | 47.36 |  | 5160.6383 | 5160.6341 | 0.82 |
| 124 | Neu5Ac_4_HexNAc_6_Hex_8_Fuc_2_Red-HexNAc_1_ | 1 | 69.03 |  | 5203.6309 | 5203.6287 | 0.44 |
| 125 | Neu5Ac_4_HexNAc_6_Hex_9_Fuc_1_Red-HexNAc_1_ | 1 | 68.57 |  | 5233.6412 | 5233.6393 | 0.38 |
| 126 | Neu5Ac_4_HexNAc_6_Hex_9_Fuc_2_Red-HexNAc_1_ | 1 | 62.20 |  | 5407.7311 | 5407.7285 | 0.50 |
| 127 | Neu5Ac_4_HexNAc_7_Hex_8_Fuc_2_Red-HexNAc_1_ | 1 | 66.82 |  | 5448.7543 | 5448.7550 | 0.12 |
| 128 | Neu5Ac_4_HexNAc_7_Hex_9_Fuc_1_Red-HexNAc_1_ | 1 | 68.07 |  | 5478.7664 | 5478.7656 | 0.16 |
| 129 | Neu5Ac_2_HexNAc_10_Hex_9_Fuc_1_Red-HexNAc_1_ | 2 | 46.72, 47.73 |  | 5491.7970 | 5491.7972 | 0.03 |
| 130 | Neu5Ac_3_HexNAc_7_Hex_10_Fuc_2_Red-HexNAc_1_ | 1 | 68.12 |  | 5495.7868 | 5495.7809 | 1.09 |
| 131 | Neu5Gc_1_Neu5Ac_3_HexNAc_7_Hex_9_Fuc_1_Red-HexNAc_1_ | 1 | 67.81 |  | 5508.7761 | 5508.7762 | 0.01 |
| 132 | Neu5Ac_4_HexNAc_8_Hex_10_Fuc_1_Red-HexNAc_1_ | 1 | 68.66 |  | 5927.9920 | 5927.9917 | 0.07 |
| **Phosphorylated *N*-glycans** | | | | | | | |
| 133 | HexNAc_1_P-Hex_1_Hex_3_Red-HexNAc_1_ | 1 | 51.69 |  | 1462.7076 | 1462.7069 | 0.50 |
| 134 | HexNAc_1_P-Hex_1_Hex_4_Red-HexNAc_1_ | 1 | 56.62 |  | 1666.8076 | 1666.8067 | 0.58 |
| 135 | HexNAc_1_P-Hex_1_Hex_4_Fuc_1_Red-HexNAc_1_ | 1 | 59.49 |  | 1840.8960 | 1840.8959 | 0.08 |
| 136 | HexNAc_1_P-Hex_1_Hex_5_Red-HexNAc_1_ | 1 | 59.04 |  | 1870.9077 | 1870.9064 | 0.69 |
| 137 | HexNAc_1_P-Hex_1_Hex_5_Fuc_1_Red-HexNAc_1_ | 1 | 61.70 |  | 2044.9965 | 2044.9956 | 0.43 |
| 138 | Neu5Ac_1_HexNAc_2_P-Hex_1_Hex_5_Fuc_1_Red-HexNAc_1_ | 1 | 62.04 |  | 2651.2954 | 2651.2956 | 0.08 |
| 139 | Neu5Ac_1_HexNAc_2_P-Hex_1_Hex_6_Fuc_1_Red-HexNAc_1_ | 1 | 64.78 |  | 2855.3912 | 2855.3954 | 1.46 |
| 140 | Neu5Ac_2_HexNAc_3_P-Hex_1_Hex_7_Fuc_1_Red-HexNAc_1_ | 1 | 66.14 |  | 3665.7973 | 3665.7952 | 0.59 |

Table S5. The identified glycopeptides by chymotrypsin digestion and comparison with permethylated N-glycan analysis.

| Peptide sequence | (LL)EAKEAE**N**(24)ITTGCcAEHCcSL | | | | | | | | | | | | |
| --- | --- | --- | --- | --- | --- | --- | --- | --- | --- | --- | --- | --- | --- |
| N-glycan structure |  |  |  |  |  |  |  |  |  |  |  |  |  |
| Glycopeptide | ***√*** | ***√*** | ***√*** | ***√*** | ***√*** | ***√*** | ***√*** | ***√*** | ***√*** | ***√*** | ***√*** | ***√*** | ***√*** |
| N-glycan | ***√*** | ***√*** | ***√*** | ***×*** | ***√*** | ***×*** | ***×*** | ***√*** | ***√*** | ***√*** | ***×*** | ***√*** | ***√*** |
|  | ***√*** | ***√*** | ***√*** | ***√*** |  |  |  |  |  |  |  |  |  |
|  | ***√*** | ***√*** | ***×*** | ***√*** |  |  |  |  |  |  |  |  |  |
| Peptide sequence | NE**N**(38)ITVPDTKVNF(Y) | | | | | | | | | | | | |
| N-glycan structure |  |  |  |  |  |  |  |  |  |  |  |  |  |
| Glycopeptide | ***√*** | ***√*** | ***√*** | ***√*** | ***√*** | ***√*** | ***√*** | ***√*** | ***√*** | ***√*** | ***√*** | ***√*** | ***√*** |
| N-glycan | ***√*** | ***×*** | ***√*** | ***√*** | ***×*** | ***√*** | ***√*** | ***×*** | ***√*** | ***√*** | ***√*** | ***√*** | ***√*** |
| Peptide sequence | (L)V**N**(83)SSQPW(EPL) | | | | | | | | | | | | |
| N-glycan structure |  |  |  |  |  |  |  |  |  |  |  |  |  |
| Glycopeptide | ***√*** | ***√*** | ***√*** | ***√*** | ***√*** | ***√*** | ***√*** | ***√*** | ***√*** | ***√*** | ***√*** | ***√*** | ***√*** |
| N-glycan | ***√*** | ***√*** | ***×*** | ***√*** | ***√*** | ***×*** | ***√*** | ***√*** | ***×*** | ***×*** | ***×*** | ***√*** | ***×*** |
|  | ***√*** | ***√*** | ***√*** | ***√*** | ***√*** | ***√*** | ***√*** | ***√*** | ***√*** | ***√*** | ***√*** | ***√*** | ***√*** |
|  | ***√*** | ***×*** | ***×*** | ***×*** | ***√*** | ***√*** | ***×*** | ***√*** | ***×*** | ***×*** | ***√*** | ***×*** | ***×*** |
|  | ***√*** | ***√*** | ***√*** | ***√*** | ***√*** | ***√*** | ***√*** | ***√*** | ***√*** | ***√*** | ***√*** | ***√*** | ***√*** |
|  | ***√*** | ***×*** | ***√*** | ***√*** | ***√*** | ***√*** | ***×*** | ***√*** | ***√*** | ***√*** | ***√*** | ***√*** | ***√*** |
|  | ***√*** | ***√*** | ***√*** | ***√*** | ***√*** | ***√*** | ***√*** | ***√*** | ***√*** | ***√*** |  |  |  |
|  | ***×*** | ***×*** | ***√*** | ***×*** | ***×*** | ***×*** | ***√*** | ***×*** | ***√*** | ***√*** |  |  |  |

Figure S1. The newly designed Python script for the identification of permethylated N-glycans, considering trimannosylchitobiose core.

Figure S2. The additionally designed Python script, especially for the identification of permethylated N-glycans containing phosphorylated Man.

Figure S3. The characterization of Fuc_1_Red-HexNAc_1_, by EIC and MS2 fragmentation from N-glycans of EPO.

Figure S4. The identified peptides of EPO-3 based on bottom-up analysis after trypsin and chymotrypsin digestion by pFind software, with the known modifications.

Figure S5. The characterization of GAQKEAISPPDAAS(126)AAPL at MS2 level from chymotryptic digestion.

Figure S6. The comparison of N-glycan compositions identified from EPO-3 at different levels, including glycopeptide analysis using pGlyco software and permethylated N-glycan analysis by the Python scripts.
